# Essential and virulence-related protein interactions of pathogens revealed through deep learning

**DOI:** 10.1101/2024.04.12.589144

**Authors:** Ian R. Humphreys, Jing Zhang, Minkyung Baek, Yaxi Wang, Aditya Krishnakumar, Jimin Pei, Ivan Anishchenko, Catherine A. Tower, Blake A. Jackson, Thulasi Warrier, Deborah T. Hung, S. Brook Peterson, Joseph D. Mougous, Qian Cong, David Baker

## Abstract

Identification of bacterial protein–protein interactions and predicting the structures of the complexes could aid in the understanding of pathogenicity mechanisms and developing treatments for infectious diseases. Here, we developed a deep learning-based pipeline that leverages residue-residue coevolution and protein structure prediction to systematically identify and structurally characterize protein-protein interactions at the proteome-wide scale. Using this pipeline, we searched through 78 million pairs of proteins across 19 human bacterial pathogens and identified 1923 confidently predicted complexes involving essential genes and 256 involving virulence factors. Many of these complexes were not previously known; we experimentally tested 12 such predictions, and half of them were validated. The predicted interactions span core metabolic and virulence pathways ranging from post-transcriptional modification to acid neutralization to outer membrane machinery and should contribute to our understanding of the biology of these important pathogens and the design of drugs to combat them.

## Introduction

Understanding the biology of pathogenic bacteria is important for human health and therapeutics. Protein-protein interactions (PPIs) are central to biological processes, but many interactions remain unknown, especially for non-model organisms. High-throughput experiments such as the two-hybrid screen and affinity purification coupled with mass spectrometry (AP/MS) have been used to identify PPIs in a variety of organisms (*1–3*). However, such methods can fail to reveal transient interactions and be plagued by non-specific interactions in non-physiological conditions, which result in discrepancies between experiments along with high false-positive and false-negative rates (*4, 5*). Interacting proteins often co-evolve, and hence amino acid coevolution can be exploited to assess the likelihood that two proteins interact with each other. Coevolutionary information between proteins extracted from paired multiple sequence alignments (pMSAs) of orthologous proteins (*6–8*) has been used to systematically identify PPIs in prokaryotes at an accuracy that rivals experimental screens (*7*). Supplementing coevolution with deep learning (DL) based structure prediction methods has further increased the accuracy of PPI prediction, enabling large-scale prediction of PPIs in yeast (*9*) and humans (*10, 11*).

We set out to systematically identify and structurally characterize PPIs in pathogenic bacteria. We selected 19 bacterial pathogens (table S1) that span 6 phyla and are the leading causes of pathogen-associated deaths in humans (12). These organisms are associated with infections in skin (*Staphylococcus aureus*), gastrointestinal tract (*Clostridioides difficile, Helicobacter pylori, Listeria monocytogenes*, and *Salmonella typhimurium*), respiratory system (*Legionella pneumophila, Mycobacterium tuberculosis, Pseudomonas aeruginosa, and Streptococcus pneumoniae*), urinary and genital tracts (*Chlamydia trachomatis* and *Mycoplasma genitalium*), and the plague (*Yersinia pestis*).

For most of the selected organisms, large-scale experimental screens have been conducted to identify essential genes (EGs), which have been collected in the Database of Essential Genes (DEG) (*12*). Many virulence factors (VFs) have been experimentally characterized in these infectious bacteria, and these results are summarized in the Virulence Factor DataBase (VFDB) (*13*). We primarily focused on EGs and VFs in our PPI study because the former provides targets for drug development to inhibit essential cellular functions and treat infectious diseases, while the latter may explain molecular mechanisms of pathogenicity. Comparative analysis showed significant overlap in the sets of EGs between different pathogens, but each pathogen still harbors ∼100 unique EGs (table S2). In contrast, VFs differ considerably between species, suggesting a diversity of virulence mechanisms exploited by different pathogens, which we attempt to capture with our set of phylogenetically diverse species (table S2).

### Computational pipeline for proteome-wide PPI identification

To screen through hundreds of millions of protein pairs for PPIs, we first sought to increase the computational efficiency of PPI identification without compromising accuracy. We previously developed a 2-track RoseTTAFold (RF 2-track) network that is a simplified version of RoseTTAFold (*14*). Although RF 2-track was not trained to model protein complexes or distinguish interacting from non-interacting proteins, residue-residue distograms produced by this network enable the detection of PPIs on a proteome-wide scale at an accuracy that far exceeds statistical analysis of coevolution between proteins (*9*). Similarly, we and others have used AlphaFold (AF) (*15*) to evaluate interactions identified in lower accuracy large-scale screens (*9–11, 16, 17*); the computational cost of AF prohibits its application on a protome-wide scale. AF-multimer (*18*) was trained to model 3D structures of known protein complexes, and consequently, it tends to predict PPIs between non-interacting pairs, displaying a worse performance in distinguishing true PPIs from random pairs than AF (**Fig. 1B** top).

**Figure 1:**
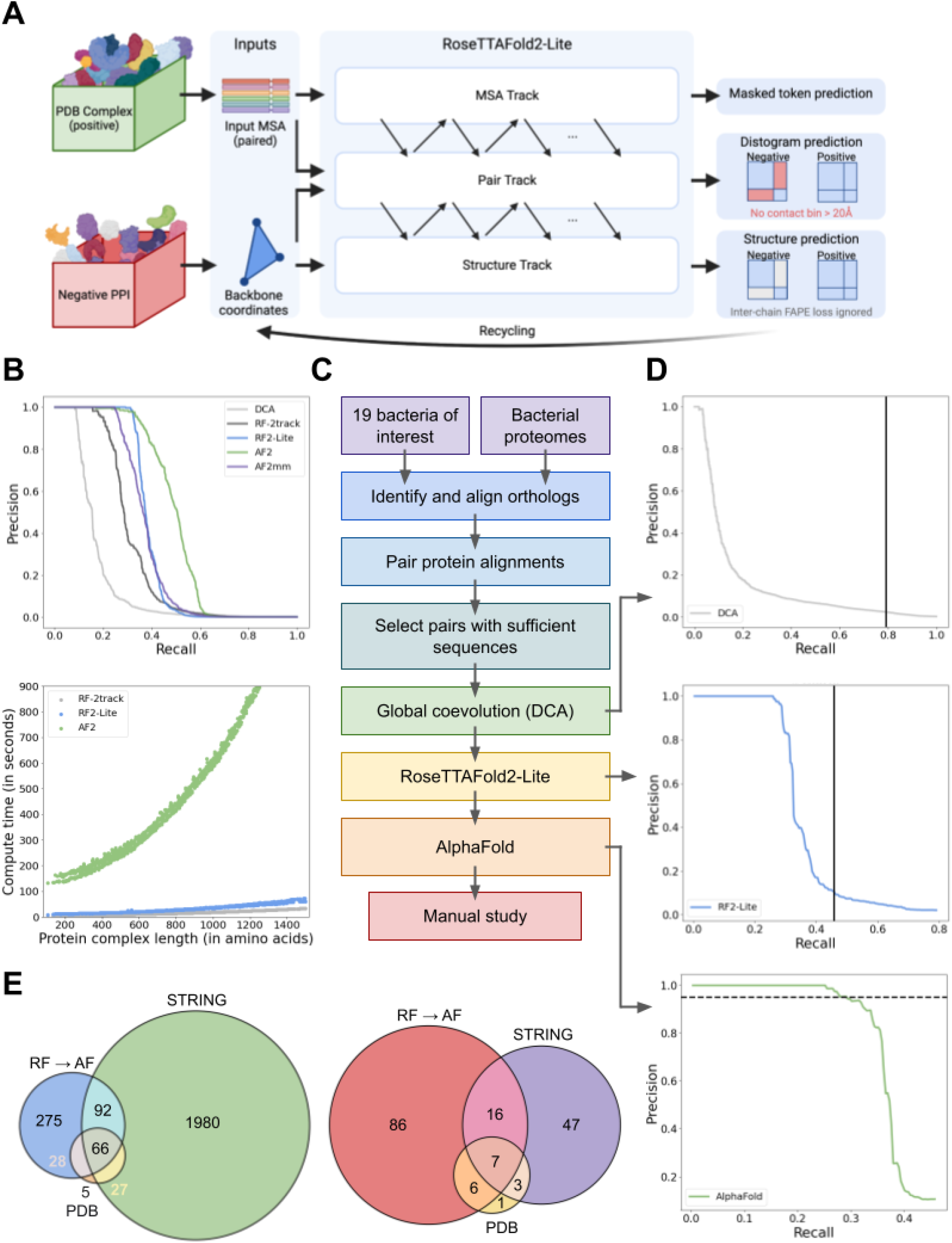
Protein-protein interaction identification by coevolution and deep learning methods. **(A)** Overview of the RF2-Lite network architecture. (**B**) Benchmark performance of PPI prediction methods. Top: precision and recall curves of DCA (grey), RF 2-track (black), RF2-Lite (blue), AF (green), and AF-multimer (purple) in distinguishing true PPIs from random protein pairs. For different methods, we used the same pMSAs generated by our bioinformatic pipeline described. We applied each method on a benchmark set of 1000 randomly selected positive control pairs and 10,000 negative control pairs. These control sets were assembled based on information about the 19 pathogens in STRING. The precision and recall curve for this benchmark is in fig. S6A. Because the real signal-to-noise ratio for the PPI screen is on the order of 1:1000 (*1*), to reflect the impact of a much larger set of non-interacting pairs, we upsampled the negative control set to 1,000,000 by randomly sampling 100 “pseudo” interacting probabilities from the Gaussian distribution around each real interacting probability we obtained for the negative controls with a standard deviation of 0.1. Bottom: runtime comparison of different methods. **(C)** Schematic overview of our PPI screen pipeline. **(D)** Precision and recall curves at different stages in the pipeline. Top: precision and recall curve of DCA on PPI prediction; solid black vertical line represents the recall cutoff in this stage. Middle: precision and recall of our RF2-Lite screen procedure on the ‘pilot set’; solid black vertical line indicates the recall cutoff at this stage. Bottom: precision and recall of our AF screen procedure on the ‘pilot set’; dashed horizontal line shows the precision cutoff, i.e., 0.95. **(E)** Summary of predicted PPIs for the “pilot set” that focuses on EGs and VFs. Left: interactions between interacting EGs in the ‘pilot set’ based on different evidence: blue, green, and orange circles represent our predicted pairs, functional interactions according to STRING (total score ≥ 900 and experimental score ≥ 400), and interacting pairs according to PDB (BLAST hit to complex in PDB e-value ≤ 0.00001, sequence identity ≥ 50% and coverage ≥ 50%), respectively. Right: PPIs involving VFs in the ‘pilot set’ supported by difference evidence: red, purple, and yellow circles represent our predictions, pairs according to STRING, and pairs according to PDB.

We hypothesized that a dedicated lighter-weight network trained on both interacting and non-interacting protein pairs that balances accuracy with speed could assist proteome-wide PPI screens. We revised the original RF network by introducing architectural improvements to increase accuracy while reducing the number of layers to enable the rapid computation necessary for large-scale screens (**Fig. 1A**). We trained this network using a combination of (1) monomeric protein structures from Protein Data Bank (PDB), (2) AF models of UniRef50 sequences, (3) pairwise protein complex structures extracted from PDB, and (4) random non-interacting protein pairs. The four types of training data were mixed at a ratio of 1:3:2:2 (table S3). The model was trained using the masked language model (MLM) loss, distogram (dist) prediction loss, frame aligned point error (FAPE) loss, accuracy estimation loss, bond geometry loss, and van der Waals (vdW) energy loss. For the negative interaction examples, we ignored the inter-chain region for FAPE calculation and required the network to predict the distogram to be in the “non-interacting bin” for the inter-chain region. We designate the resulting network as RoseTTAFold2-Lite (RF2-Lite) as it resembles the RoseTTAFold2 architecture but has many fewer parameters (*19*). RF2-Lite has improved performance in distinguishing true PPIs over the previous RF 2-track: at the same precision, the recall for true PPIs by RF2-Lite is in between RF 2-track and AF (**Fig. 1B** top; fig. S6-S7). Despite this increase in accuracy, RF2-Lite’s speed is still comparable to RF 2-track, and it requires about 20-fold less compute time than AF (**Fig. 1B** bottom).

We combined direct coupling analysis (DCA) (*20*), RF2-Lite, and AF (**Fig. 1C**) to identify and model interacting proteins and applied this pipeline to the 19 human pathogens listed in table S2. To monitor the performance of our pipeline, we assembled a set of positive controls and a ∼700-fold larger negative set based on information from the STRING database. The ratio between the positive and negative controls was based on the assumption that each protein, on average, possesses 5 interacting partners out of all other proteins from a species. The positive controls consist of protein pairs from these 19 pathogens with a STRING experimental score > 600 and a STRING combination score > 990. Pairs of proteins from the ribosome or NADH-quinone oxidoreductase complex were excluded because nearly all such pairs receive high STRING scores but most do not interact directly. The negative controls are random pairs with no evidence supporting interaction in STRING.

We constructed a database of 44,871 representative bacterial proteomes/genomes (one per species) obtained from NCBI and used the reciprocal best hit criteria (*21*) to identify an orthologue for every protein in each proteome (fig. S1). We aligned these orthologous sequences (*22, 23*), and for each protein pair in each of the 19 pathogens (fig. S2), we concatenated their MSAs by connecting sequences of the same species to generate pMSAs (fig. S3). We removed proteins whose monomeric structure could not be confidently modeled by AF (average pLDDT < 50 in AFDB) and filtered the pMSAs based on their depth and quality (figs. S4,S5): of the total 140.2 million protein pairs, we selected 77.9 million (56%) with higher monomer structure and MSA quality.

We assessed the residue-residue coevolution for the selected pairs using DCA, and found that the 7.7 million (10%) high-scoring protein pairs by DCA contained 79% of the positive controls (**Fig. 1D** top). Among these 7.7 million pairs, we initially focused on a “pilot set” of 0.14 million pairs involving at least one VF (according to VFDB) and 0.83 million pairs of EGs (according to DEG). We removed redundancy in this set by clustering proteins from the 19 species into orthologous groups using OrthoMCL (*24*). If the orthologs of a protein pair were present in multiple species, we selected only one pair with the highest DCA score, resulting in a total of 457,310 representative PPI candidates.

We used RF2-Lite to identify confident PPIs from the “pilot set” and observed that we could achieve a recall of 28% at a precision of 95% when an RF2-Lite contact probability cutoff of 0.74 was used (**Fig. 1D** middle). We investigated whether using a loose RF2-Lite cutoff (contact probability 0.05) to select candidate PPIs (46,609, around 10% selected) for AF could improve recall. The RF2-Lite → AF pipeline only improved the recall to 29% at 95% precision (**Fig. 1D** bottom) at the cost of using 3 times more computer resources than simply relying on RF2-Lite to detect PPIs (table S4). Thus, the contribution of AF in distinguishing true PPIs from random pairs is limited, but it remains essential for obtaining high-quality 3D structures for the predicted protein complexes.

The successive use of DCA (selecting top 10%), RF2-Lite (cutoff: 0.05), and AF (cutoff: 0.9) collectively reduced the total number of random pairs by nearly 10,000 fold, resulting in 562 highly confident predictions from the “pilot set”. The identified binary protein complexes include 461 protein complexes involving EGs (**Fig. 1E** left) and 115 involving VFs (**Fig. 1E** right). Further investigation of these interactions may be useful for understanding the mechanisms of pathogenicity and developing disease prevention and treatment strategies. The vast majority (19%) of predicted protein complexes from the “pilot set” did not have experimental 3D structures in PDB (BLAST e-value ≤ 0.00001, identity ≥ 50%, and coverage ≥ 50% for both proteins), and half do not have confident experimental support according to STRING (*25*) (table S5).

To gain more structural and functional insights into these pathogens, we applied the RF2-Lite → AF pipeline to an additional 3.82 million pairs involving essential proteins and biological processes of therapeutic interest, such as the outer membrane machinery (table S6). This search resulted in an additional 3,051 predicted PPIs. Based on our positive and negative controls, PPIs identified by our pipeline have a predicted precision of 95%. This high level of precision is based on the assumption that each protein is expected to directly interact with 5 other proteins. However, if the signal-to-noise ratio is much lower, e.g., the average number of direct interacting partners for each protein is 1, the estimated precision falls to 80%. Inspection of the predicted PPIs revealed a small number of proteins (in particular ferredoxin and rubredoxin) with predicted interactions between many random proteins, likely constituting small false positive hubs. We removed 405 PPIs involving such potential false positive hubs before deposition to ModelArchive.

It is difficult to cover even a small fraction of the biological insights that can be revealed from these 3D structures of protein complexes in one paper. In the following sections, we first describe experimental validation for a subset of predictions and then highlight examples that illustrate some of the biological insights revealed by the identification of putative PPIs and computational modeling of protein complexes.

### Experimental validation

To corroborate our benchmarking analyses, which suggest that our predicted interactions should be quite accurate, we selected two sets of predicted interactions for experimental characterization. We biased these selections towards PPIs with no prior experimental evidence or strong functional associations because validating such interactions could provide new biological insights. The first set (table S7) was selected based on statistical methods (GREMLIN) for PPI detection, prior to the development and application of the DL methods. This set was used to probe the accuracy of statistical (DCA and GREMLIN (*26*)) versus DL methods for PPI detection. The second set (table S8) was selected from our final set of predicted 3,613 PPIs, with a goal of evaluating the accuracy of our current entire pipeline.

We selected the first dataset using the following criteria: (1) at least 20 kb apart (with a minimum of 20 intervening genes), (2) not having homologous complexes in the PDB, (3) not predicted to have the same molecular function, (4) not annotated as part of the same biological pathway, and (5) not strongly supported by STRING (combined score < 800). All eleven pairs show strong coevolution according to DCA and GREMLIN, but five pairs were not predicted to interact by RF2-Lite or AF (fig. S11). A bacterial two-hybrid (B2H) system (*27*) coupled with a quantitative β-Galactosidase assay (*28*) was employed to measure interactions for these eleven pairs (fig. S12).

Despite the strong support by DCA and GREMLIN, the five pairs not predicted to interact by RF2-Lite or AF did not show evidence of interaction using the B2H assay (fig. S11). Among the six pairs supported by RF2-Lite or AF, significant reporter activation indicative of interaction was detected for two: one is between iron-sulfur cluster binding protein lpg2881 (Uniprot: Q5ZRK0) and uncharacterized protein lpg0371 (Uniprot: Q5ZYK1) from *L. pneumophila*; another is between ribosomal silencing factor RsfS (PA4005; Uniprot: Q9HX22) and PhoH-like protein domain-containing protein YbeZ (PA3981; Uniprot: Q9HX38) from *P. aeruginosa* (**Fig. 2A**). For one additional pair, nucleoid-associated protein lmo2703 (Uniprot: Q8Y3X6) and signal recognition particle protein Ffh (Uniprot: Q8Y695) from *L. monocytogenes*, we were unable to assess the interaction experimentally due to false-positive reporter activation when only one protein was expressed (fig. S12). The remaining three pairs failed to generate a positive reporter signal; however, false negative results from B2H assays do not necessarily rule out the existence of a genuine interaction due to possible failures in protein expression and folding of the fusion proteins, and lack of sensitivity of the screen to weak and transient interactions.

**Figure 2:**
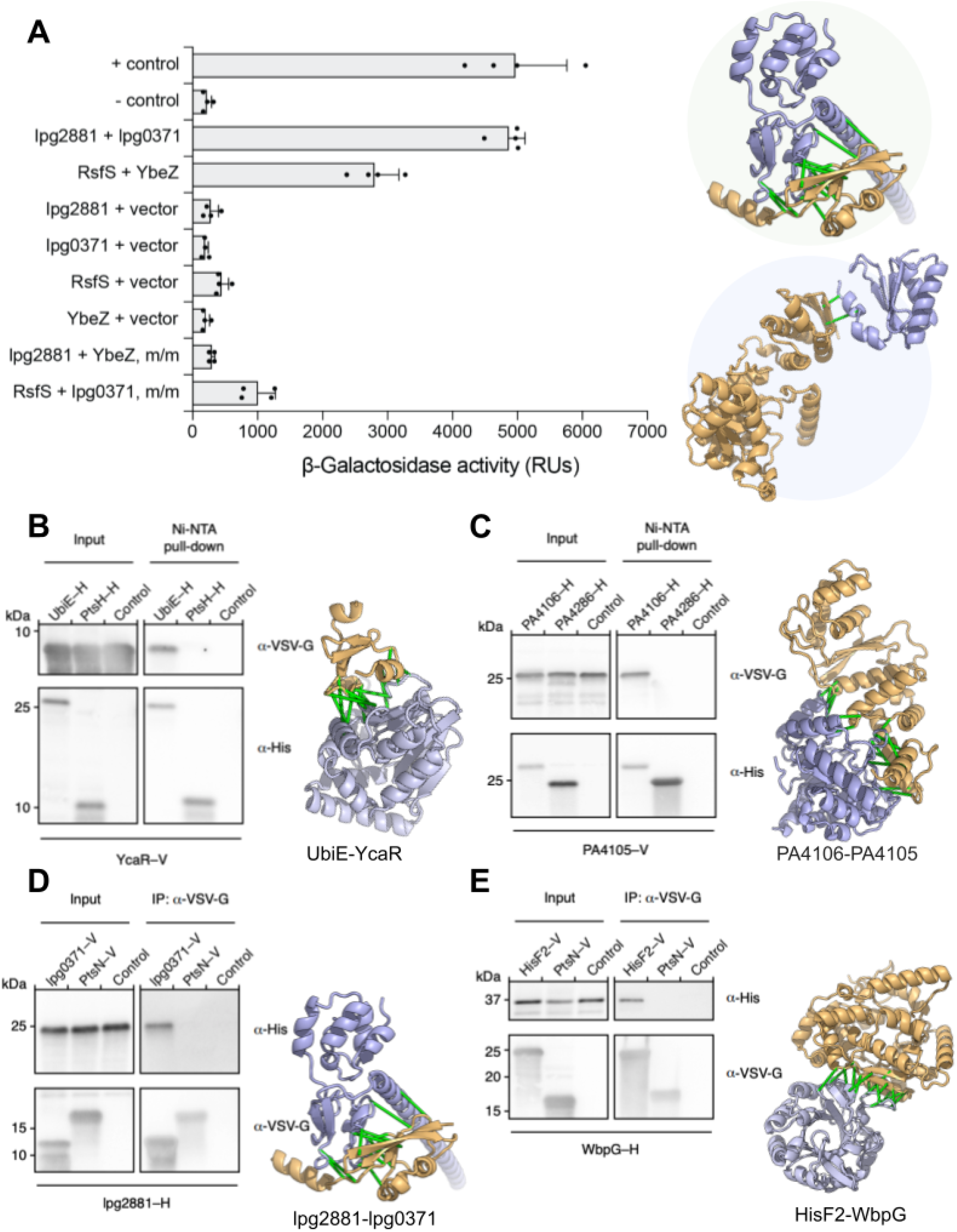
Experimental validation of selected protein-protein interactions. **(A)** Interactions assessed by B2H that measures β−galactosidase activity resulting from activation of the *lacZ* reporter gene due to the interaction between two tested proteins that are fused to two domains of a transcription activator. *E. coli* expressing T25-zip and T18-zip fusion proteins was used as a positive control (+ control), and *E. coli* harboring empty T25 and T18 plasmids was used as a negative control (-control). m/m = mix-and-match control. RUs (relative units) = Luminescence / OD_600_ per hour. Error bars indicate ±s.d. (*n* = 2 biological replicates each with 2 technical replicates). Computed models of experimentally validated PPIs (lpg2881 + lpg0371, and RsfS + YbeZ) are shown on the right: top, iron-sulfur cluster binding protein lpg2881 (Q5ZRK0) and uncharacterized protein lpg0371 (Q5ZYK1) from *L. pneumophila*; bottom, ribosomal silencing factor RsfS (Q9HX22) and PhoH-like protein domain-containing protein YbeZ (Q9HX38) from *P. aeruginosa*. **(B-E)** Interactions validated by co-immunoprecipitation (Co-IP)/pull-down. Predicted interacting partners in each PPI pair are heterologously expressed and tagged (–H, hexahistidine; –V, VSV-G epitope). A random bait protein was included as a negative control for each experiment. Control lanes correspond to samples with prey proteins and beads added without any bait proteins. Each positive interaction is supported by two independent Co-IP/pull-down experiments. **(B)** Ubiquinone biosynthesis C-methyltransferase UbiE (P0A887) and protein of unknown function YcaR (P0AAZ7) from *E. coli*. **(C)** Uncharacterized protein PA4106 (Q9HWS2) and a putative transcriptional factor PA4105 (Q9HWS3) from *P. aeruginosa*. **(D)** lpg2881 and lpg0371 from *L. pneumophila*, a pair that is tested positive by B2H as well. **(E)** Putative imidazole glycerol phosphate synthase subunit hisF2 (P72139) and LPS biosynthesis protein WbpG (Q9HZ78) from *P. aeruginosa*. In all the panels, connecting green bars are between representative residue-residue contacts at the interfaces predicted from the summed AlphaFold probability for distance bins below 12Å.

For both PPIs validated by our B2H assays, there are no published data directly supporting functional or physical interactions between the two proteins. However, in both cases, existing evidence indirectly suggests that the interactions could be biologically significant. The pair of proteins from *L. pneumophila* (lpg2881-lpg0371; Q5ZRK0-Q5ZYK1) are homologous to proteins of the Rnf electron transport complex (RnfB with 53% sequence identity and RnfH with 36% sequence identity, respectively). The function of these proteins in *L. pneumophila* is unclear because this species appears to lack the other components of the complex, and one of the proteins, lpg0371, also shares homology with the antitoxin component of the RatAB toxin-antitoxin module. However, in species that encode the complete Rnf complex, RnfB and RnfH directly interact (*29*). The interacting pair from *P. aeruginosa* consists of the ribosomal silencing factor RsfS and the PhoH-like protein domain-containing protein YbeZ. Under nutrient depletion or during stationary phase growth, RsfS binds to ribosomal protein L14, ultimately preventing the association of the 30S and 50S ribosomal subunits and repressing translation (*30*). This facilitates adaptation to low nutrient conditions and promotes survival during the stationary phase. The function of YbeZ is less well-characterized, but it interacts with the RNase YbeY, and both proteins are required for processing and maturation of the 16S rRNA (*31*). Our finding that YbeZ and RsfS interact suggests that the regulation of ribosome assembly and ribosome subunit processing may be linked in *P. aeruginosa*.

The second validation set consists of six protein pairs (table S8). Similarly to the first experimentally tested set, none of the PPI candidates in this second set have homologous protein complexes in the PDB. Other characteristics typical of interacting protein pairs varied across the candidates: most interactions were not supported by STRING (one pair STRING > 600), and half of the candidates are not genomic neighbors (> 100 genes apart). Finally, we opted to focus on proteins consisting primarily of globular domains (percentage of residues from non-globular domains < 20%), as such proteins are more amenable to heterologous expression-based assays, and we included one interacting pair we had validated by B2H (lpg2881-lpg0371).

Using co-immunoprecipitation (Co-IP) assays, we detected an interaction between four of the six pairs in the second test set. As expected, this included the pair we had previously validated by B2H, Q5ZRK0-Q5ZYK1, a distally encoded pair from *E. coli* and two proximally encoded pairs from *P. aeruginosa*. These validated PPIs span the spectrum from those for which no previous data hints at a functional relationship to pairs believed to participate in the same pathway but for which no interaction was previously predicted. The interacting pair from *E. coli*, the carbon methyltransferase C-methyltransferase UbiE (P0A887) and the uncharacterized protein YcaR (P0AAZ7) fall into the former category. UbiE catalyzes a carbon-methyl transfer reaction in the biosynthesis of ubiquinone (coenzyme Q) and menaquinone (vitamin K2) (*32*), while YcaR is a small protein detected as differentially expressed in multiple proteomics studies but to which no function has been assigned (*33, 34*). Our discovery of an interaction between these proteins suggests a number of possible hypotheses for YcaR function that can now be interrogated. One of the validated interacting pairs from *P. aeruginosa*, PA4105-PA4106 (Q9HWS3-Q9HWS2), consists of two uncharacterized proteins with no clear homologs of known functions based on primary sequence comparisons. However, a FoldSeek search (*35*) revealed structural similarity between these proteins and TglI and TglH from *P. syringae pv. maculicola* (*P. syringae*), respectively. TglI and TglH form a complex that catalyzes the removal of cysteine β-methylene (β-CH2) from TglA-Cys, a step in the biosynthesis of the natural product 3-thiaglutamate (3-thiaGlu) (*36, 37*). Our detection of an interaction between Q9HWS3 and Q9HWS2 suggests these proteins may perform a similar reaction. The last validated interaction is between two VFs of P. aeruginosa: HisF2 (P72139) and WbpG (Q9HZ78). WbpG is an amidotransferase that is essential in B-band LPS biosynthesis, and HisF2 is a predicted imidazole glycerol phosphate synthase subunit. LPS has been linked to the virulence of *P. aeruginosa*, allowing this pathogen to resist serum killing and phagocytosis and contributing to its invasion into host cells. It was previously proposed that HisF2, together with HisH2, delivers ammonia to WbpG (*38*), a hypothesis our interaction finding supports. Surprisingly, the PtsH-PtsN (Q9HVV2-Q9HVV4) pair with the highest support by STRING (score = 959) failed to generate a positive Co-IP signal (fig. S13). PtsH is a histidine-phosphorylatable phosphocarrier protein encoded adjacent to PtsN, a nitrogen regulatory protein with a phosphotransferase component. We hypothesize that the interaction between these proteins may be transient, and thus difficult to detect by co-IP.

More broadly, these experimental data support the *in silico* benchmark in suggesting that the DL methods have greater accuracy than statistical methods in PPI discovery. The examples above show that DL methods can identify additional components for well-known biological pathways and accelerate the characterization of proteins of unknown function. In the following sections, we provide an overview of the much larger set of interactions predicted by the DL methods but not yet experimentally validated; to illustrate the insights that can be gained from these data, we provide biological context for selected interaction pairs and higher-order assemblies.

### Binary interactions

From the total set of 3613 predicted binary PPIs, 1686 (47%) have homologous complexes in PDB (BLAST e-value ≤ 0.00001 for both proteins), 1862 (52%) are supported by strong functional association according to STRING (total score ≥ 900), and 1284 (36%) are supported by both PDB and STRING; the remaining 1349 (37%, 3613 − (1686 + 1862 − 1284)) to our knowledge, are unknown PPIs. Although such previously unsupported PPIs might contain a higher fraction of false predictions, the high precision on our benchmark sets suggests a significant fraction of the new predictions are likely correct. We identify 166 putative interactions that involve uncharacterized proteins (all Pfam domains are uncharacterized), the majority of these pairs (149) include an interaction partner of known functional domains, and 131 (117 with known partners) not well described previously (STRING combined score < 900 and BLAST e-value to PDB chains > 0.00001).

1923 of the predicted PPIs include one or more EGs. Examples of predicted interactions among EGs without homologous complexes in the PDB are highlighted in **Fig. 3A-J** and table S9. In some cases, the predicted PPIs support previous findings from the literature. For example, we predict an interaction between glucose-6-phosphate 1-dehydrogenase 2 (G6PD2) and OxPP (oxidative pentose pathway) cycle protein OpcA (**Fig. 3A**). G6PD2 is an isozyme of G6PD, a member of the pentose phosphate pathway, catalyzing the oxidation of G6P to 6-phosphogluconolactone while converting NADP+ to NADPH and protecting cells from oxidative stress (*39*). OpcA has been implicated as an allosteric activator of G6PD (*40*), but, to our knowledge, the binding site remains unknown. Our predicted interface places OpcA away from the active site of G6PD, consistent with allosteric modulation of activity (fig. S19). We predict an interaction between 30S ribosomal protein S11 (rpsK), a surface-exposed ribosomal protein that forms part of the mRNA binding cleft which recognizes the Shine-Dalgarno sequence (*41, 42*), and YbeY, a highly conserved endoribonuclease which has been linked to numerous processes such as 16S rRNA maturation, 70S control, and regulation of mRNA (*43*) (**Fig. 3F**). In some bacteria, YbeY plays a key role in virulence and cell stress (*44*). Our predicted structure of S11-YbeY with an interface mediated by S11 β-strands agrees with previous work that identified S11-YbeY interaction by bacteria 2-hybrid, coimmunoprecipitation, and mutational analyses (*45*). The 3D model of the S11-YbeY complex may lend further insights into how YbeY coordinates cleavage of the rRNA precursor during 16S maturation.

**Figure 3:**
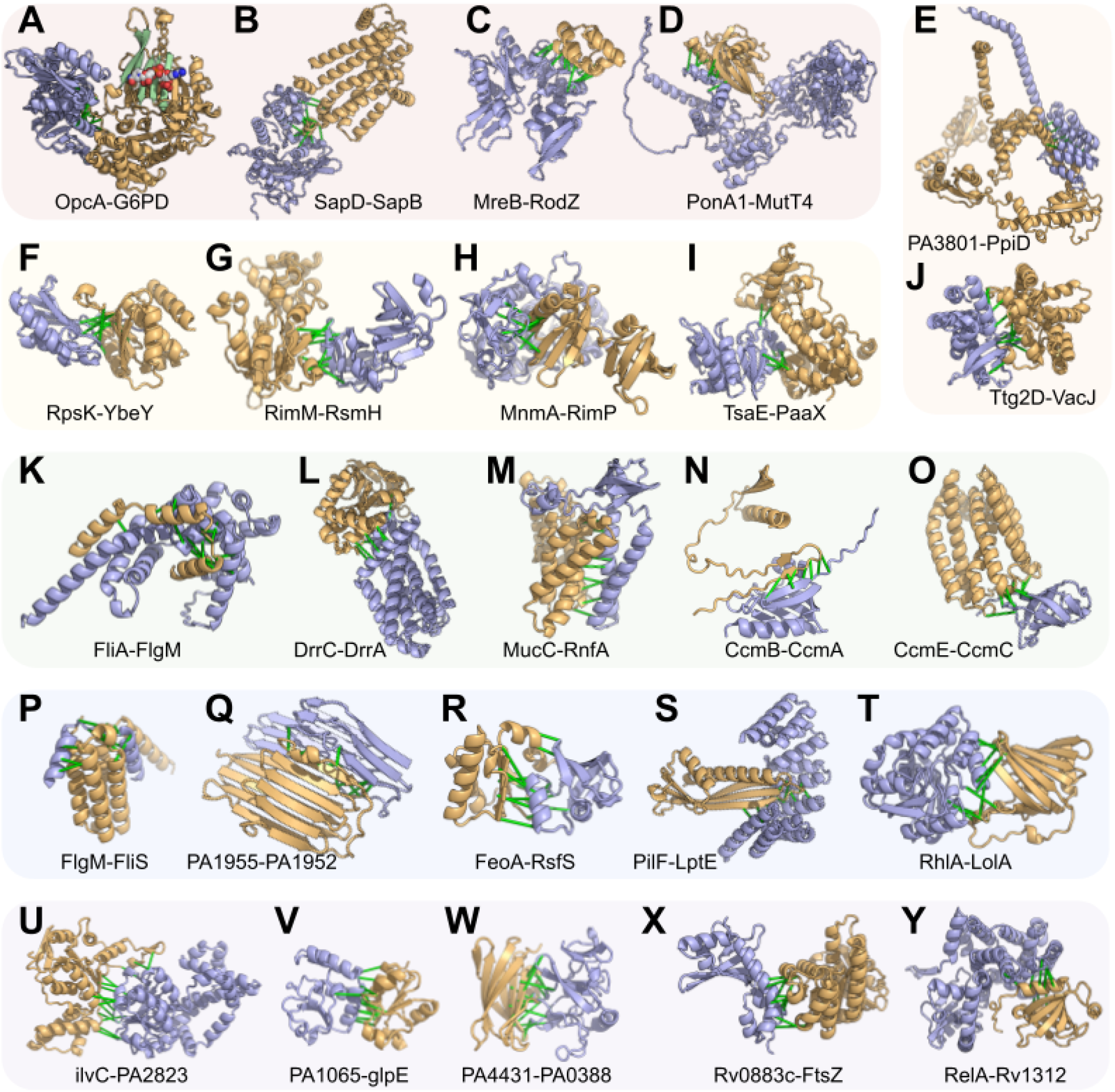
Computed models of binary protein complexes. **(A-J)** Interactions involving essential genes. **(A)** Interaction with an enzyme where the enzymatic site is highlighted in light green with an NAD moiety. **(B-D)** additional interactions involving essential genes. **(E**,**J)** Interactions involving transport pathways. **(F-I)** Transcription and translation. **(K-T)** Interactions involving virulence factors. **(U-Y)** Interactions with uncharacterized proteins. In all models, the first protein is in blue, and the second is in gold. Green bars are between representative residue-residue contacts at the interfaces predicted from the summed AlphaFold probability for distance bins below 12Å. Additional information (organisms and UniProt annotations) is in table S9.

256 of the predicted PPIs contain VFs (according to VFDB and Uniprot Keywords) that participate in pathogen colonization, nutrient acquisition, and evasion of host immunity (*46*). Secreted VFs rarely interact with endogenous proteins of a pathogen; consistent with this, we did not detect many PPIs involving VFs, and those we did identify mostly involve structural components of flagella (considered virulence factors in many bacteria (40)) and bacterial secretion systems (**Fig. 3K-T**). We also identified other interactions related to flagella function, for example, between the anti-sigma factor FlgM, a negative regulator of flagellin synthesis, and flagellar secretion chaperone (FliS) (**Fig. 3P**), an interaction supported by a previous experimental study (*47*) but without 3D structure information. Our 3D models, in agreement with previous observations (*47*), revealed that FlgM can compete with flagellin (FliC, major structural component of the flagella) for the same interface on FliS; FlgM uses its C-terminal helices to interact with FliS, which could prevent its interaction with the flagellar sigma factor FliA. The FliS-FlgM interaction might provide a negative feedback mechanism to control the expression of flagellin: when intracellular flagellin is abundant, it outcompetes FlgM in binding the anti-sigma factor FlgM, and the release of FlgM antagonizes the activity of sigma factor FliA, turning off the expression of late-stage flagellar genes, including flagellin (FliC).

We identify 149 putative interactions (**Fig. 3U-Y**) between uncharacterized proteins (according to Pfam domains) and functionally annotated binding partners such as ketol-acid reductoisomerase IlvC, thiosulfate sulfurtransferase GlpE, ubiquinol-cytochrome c reductase, cell division protein FtsZ, and bifunctional guanosine pentaphosphate [(p)ppGpp] synthase/hydrolase RelA. These predicted interaction partners provide contextual hypotheses about the function of these uncharacterized proteins, 72 of which are essential to pathogen survival, to guide further experimental studies aimed at elucidating their functions.

### Multicomponent protein complexes

In many cases, the predicted binary interactions form larger sets, suggesting the formation of higher-order assemblies. For example, in our set of 3613 predicted interactions, we found 206 trimeric protein complexes where each component is predicted to directly interact with the other two. 1545 (40%) of the predicted binary interactions involve proteins that have multiple interacting partners, which allows us to build higher-order protein complexes by concatenating the MSAs of multiple proteins and modeling them together through AF. Below we describe several selected examples of multicomponent protein complexes and the biological insights we gained from them.

#### tRNA modification and sulfur transfers in the 2-thio modification complex of E. coli

Transfer RNAs (tRNA) play critical roles in protein synthesis and are often decorated with post-transcriptional modifications that contribute to efficient protein synthesis (*48*). Wobble positions are hotspots of such modifications. In glutamate, glutamine, and lysine tRNAs, the wobble uridine is modified to 5-methylaminomethyl-2-thiouridine (mnm^5^s^2^U) by tRNA 2-thiouridine synthesizing proteins (Tus) (*49*); which include: TusA, TusB, TusC, TusD, TusE, and tRNA-specific 2-thiouridylase (MnmA). Cysteine desulfurase (IscS) is essential for 2-thio modification in *E. coli (49*). IscS transfers sulfur from cysteine to TusA, which is transferred to TusD of the TusBCD complex via TusE and subsequently to MnmA, which incorporates the sulfur into the tRNA (*49*), (*50*). The structure of the IscS-TusA dimer and sulfur transfer mediating heterohexameric complex, TusBCD, has been co-crystallized (*50, 51*), but structural details for other components of this system are poorly understood. We predicted the structures of the TusE-MnmA and TusE-TusC complexes (fig. S20) and assembled a model of the full TusBCDE heterotetramer which contextualizes the interaction of TusE with TusBCD (**Fig. 4A**; fig. S21). Our model places TusE close to TusC and TusD, with a confidently predicted TusC-TusE interface (fig. S21E) and is consistent with the hypothesis that Cys108 of TusE accepts sulfur from Cys78 of TusD (*49, 50*), but also suggests that TusC serves as a scaffold to bring TusD and TusE to close proximity. We also predict the structure of the TusE-MnmA interaction and find that TusE cannot interact with TusBCD and MnmA simultaneously due to overlap in the interfaces with MnmA and TusD (fig. S20A-F).

**Figure 4:**
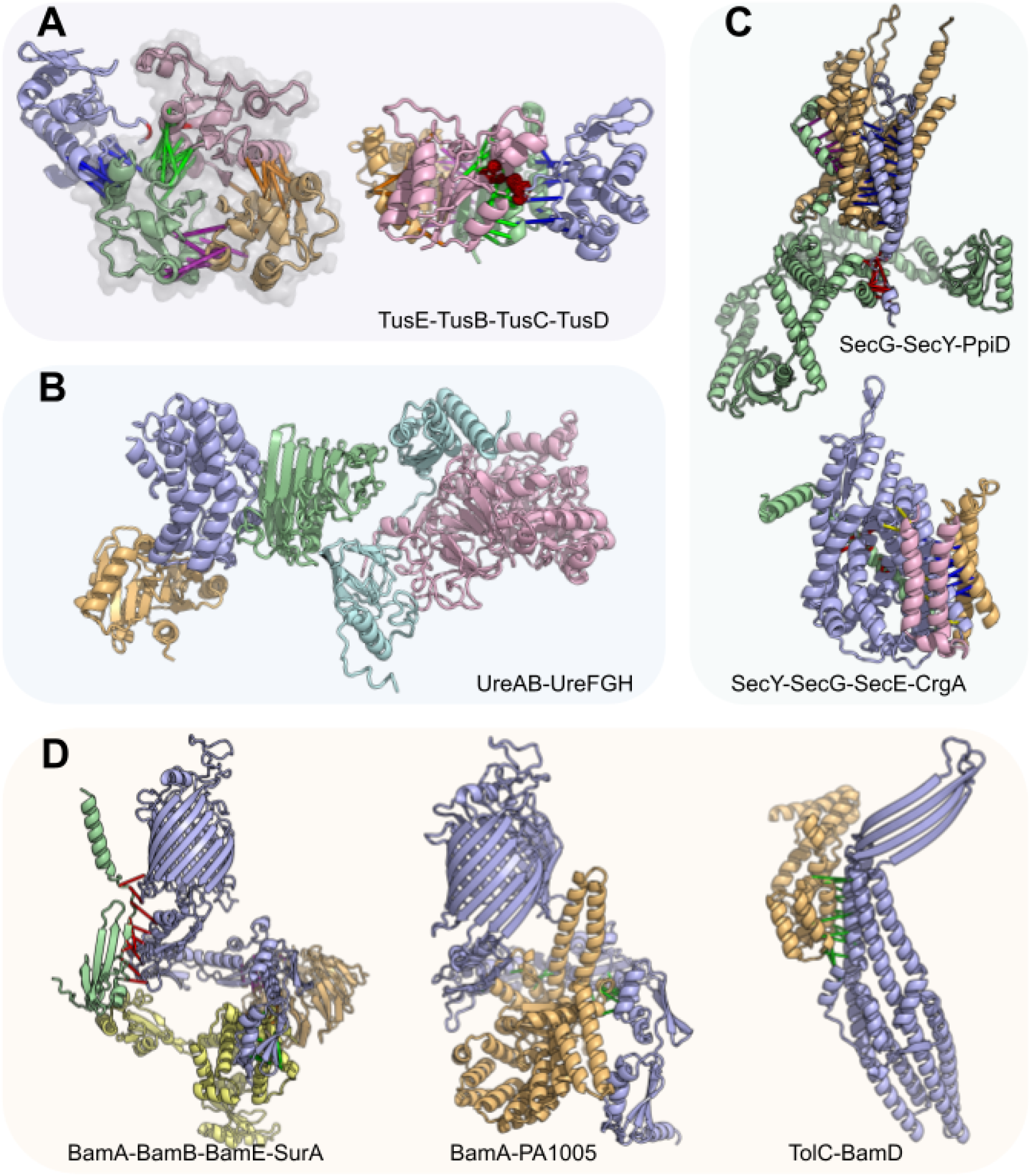
Computed models for multi-component protein complexes. **(A)** *H. pylori* tRNA 2-thiouridine synthesizing protein complex. Left: a model of the TusE(blue)-TusB(gold)-TusC(green)-TusD(pink) complex overlaid with the TusBCD PDB structure (2D1P, shown in semi-transparent grey). Right: an alternative view of this complex. **(B)** The UreAB-UreFGH complex (colored in cyan, pink, blue, gold, green, respectively) in *H. pylori* assembled through multiple subcomplexes: UreFGH, UreAB, and UreAH. **(C)** Accessory components of the Sec translocon. Top: *P. aeruginosa* SecG(blue)-SecY(gold)-PpiD(green) complex. Bottom: *M. tuberculosis* SecY(blue)-SecG(gold)-SecE(green)-CrgA(pink) complex. **(D)** Accessory components of the *P. aeruginosa and S. typhimurium* outer membrane B-barrel assembly machinery. Left: interaction between SurA (yellow) and Bam proteins (BamA: blue, BamB: gold, BamE: blue). Middle: BamA (blue) and PA1005 (gold), a putative BepA orthologue. Right: interaction between TolC (blue) and BamD (gold). In all schematics, green, red, yellow, and magenta bars connect representative residue-residue contacts at the interfaces predicted from the summed AlphaFold probability for distance bins below 12Å.

#### A two-step nickel transfer in H. pylori urease complex

Urease hydrolyses urea into ammonia and is broadly conserved in bacteria and eukaryotes. In *H. pylori*, urease neutralizes gastric acid and facilitates gut colonization (*52*), thus proteins in the urease complex are considered VFs. While most bacterial ureases have three chains (UreA, UreB, and UreC), *H. Pylori* urease has two due to the fusion of UreA and UreB orthologues (*53*). The UreAB(C) system has four accessory proteins: UreE, UreF, UreG, and UreH (*54*). We predict a UreA-UreH interaction and use it to assemble a model of a UreAB-UreFGH pentamer (**Fig. 4B**; fig. S22E). The UreAB(C) and UreFGH substructures have been determined experimentally (*53*), (*55*), and our predictions are consistent with these (fig. S22A-D). During urease maturation, UreFGH receives nickel from UreE, but how this occurs remains poorly understood. Two hypotheses are (a) that UreE transfers nickel to UreFGH complex (*56*) or (b) that upon binding GTP, UreG dissociates from UreFGH, receives nickel from UreE, and subsequently interacts with the inactive UreFH to activate the complex (*55*). Superimposing our UreE-UreG model onto the UreFGH complex shows that UreE clashes with UreF, indicating that UreE cannot directly interact with the UreFGH complex. Therefore, our observation supports the latter hypothesis wherein UreG likely receives nickel separately from UreFH (fig. S23) (*55*).

#### Interactors of the Sec translocon

The Sec translocon machinery transports proteins across the plasma membrane. The Sec translocon channel is a heterotrimeric complex composed of SecYEG, which operates in tandem with SecA, a RecA-like ATPase that moves peptides through the SecY channel in a process similar to Sec61 translocon in eukaryotes (*57*). We predict interactions between the Sec translocon and peptidyl-prolyl cis/trans isomerase D (ppiD) (**Fig. 4C** top; fig. S24), which has been identified as the most prominent interactor of SecYEG by AP/MS (*58*) and co-immunoprecipitation (*59*). In our model of the SecYG-ppiD trimer, ppiD primarily interacts with SecY through the transmembrane helices while coming close to SecG via a small loop. We also predict interactions between Sec and CrgA, a transmembrane protein and a component of the divisome (**Fig. 4C** bottom). We find that the CrgA-SecY interface occurs near the lateral gate of SecY (*60*) (fig. S25A), potentially occluding Sec translocation. We hypothesize that during bacterial division, CrgA binds Sec to regulate and recruit translocation machinery near the cell division site, this latter hypothesis is further supported by the predicted interaction between CrgA and SecE (fig. S25) and a less confident prediction of CrgA-SecG interaction that fell slightly below our cutoff.

#### Outer membrane β-barrel assembly machinery of P. aeruginosa and V. cholera

In Gram-negative bacteria, the β-barrel assembly machinery (BAM) is essential for the folding and insertion of outer membrane β-barrel proteins (*61, 62*). BAM consists of an outer membrane-spanning β-barrel, BamA, that interacts with four periplasmic lipoproteins, BamB, BamC, BamD, and BamE, to form a five-component complex (computed interactions and structures agree with known experimental data (fig. S26)) (*61–65*). This complex has recently garnered increased attention as a potential therapeutic target, especially since the discovery of Darobactin, a novel antimicrobial compound that binds along the lateral gate of BamA to inhibit outer membrane protein (OMP) biogenesis (*66, 67*).

The function of BAM is assisted by several other proteins, including the chaperone survival factor A (SurA) and periplasmic chaperone 17-kilodalton protein (Skp). SurA plays an important role in facilitating the recruitment of unfolded OMPs from the periplasm to the BAM complex (*68*). Both our BAM-SurA model and a recently published study using an orthogonal approach to ours (*69*) place SurA in the same position to simultaneously interact with BamA, BamB, and BamE (**Fig. 4D** left). Additionally, we predict an interaction between Skp and SurA (fig. S27), which in addition to their roles in maintaining the solubility of unfolded OMP proteins, may act in tandem to disassemble oligomeric OMPs that have aggregated (*70*).

We also predict an interaction between BamA and PA1005 (Uniprot: Q9I4W8) (**Fig. 4D** middle), a possible orthologue of β-barrel assembly-enhancing protease (BepA) (fig. S28). *E. coli* BepA is a periplasmic zinc-metallopeptidase with an important role in OM homeostasis and is involved in the degradation of BamA in the absence of SurA (*71*). BepA has been shown to interact with BAM (*72*), and further cross-linking experiments suggest that BepA C-terminal tetratricopeptide repeat (TPR) domain is inserted into the periplasmic region of BamA, below the β-barrel (*71*). Our computed model agrees with the proposed broad interface between BamA and BepA, provides structural details into the BamA-BepA interaction, and also suggests that when BepA is in complex with BamA, BAM is unable to assemble into its active form due to steric clashes between BepA and periplasmic Bam lipoproteins.

TolC is an outer membrane protein that homo-trimerizes to form a large outer membrane export tunnel that interacts with inner membrane translocases (*73, 74*). The catalytic β-barrel domain of BamA binds substrates along the β-barrel seam during OMP folding, and in this process, the N-terminal of the β-barrel likely swings outward (*75, 76*). The interaction between BamA and TolC has been recognized as an essential step in the assembly of TolC which occurs in a SurA-independent manner (*77, 78*). We predict an interaction between BamD and TolC (**Fig. 4D** right), which, when superimposed onto the BAM complex (fig. S29), depicts how the β-sheets of TolC interact with the N-terminal strand of the BamA β-barrel seam. Our computed model shows how TolC could be folded by the BAM complex and suggests that BamD may potentially replace SurA to stabilize or recruit TolC to BAM.

## Conclusions

RF2-Lite is a new DL network for PPI prediction that is optimized to balance the accuracy and speed necessary for large-scale applications. We integrated RF2-Lite into a pipeline for proteome-wide PPI detection and modeling. We applied this pipeline to an array of human bacterial pathogens, resulting in several thousand predicted PPIs and their 3D structure models. Over one thousand of our predictions are previously unknown, and both our benchmark and experimental validation suggest that a significant fraction of these new PPIs are likely correct and should provide novel biological insights. The 3D structure models of protein complexes generated in our study provide mechanistic details for numerous essential cellular pathways and virulence factors.

Our results demonstrate the potential of computational methods in elucidating the 3D interactome and gaining functional insights for any organism. However, there is still considerable room for improvement in reducing the false positive and false negative rates. As a consequence of the false negatives, our predictions are not comprehensive: the absence of interactions should not be overinterpreted. Although we sought to be conservative and predict only highly confident PPIs, false positives unavoidably exist in our datasets. If each protein on average interacts with only 1 partner, 80% of the predictions in our final dataset are expected to be correct based on our benchmark. Some predicted PPIs, if true, appear to be transient based on the function of proteins, and hence could be difficult to detect with experimental methods like Co-IP (without cross-linking). Based on our limited experimental validation, one should expect that ⅔ of our predicted interactions would give a positive signal in Co-IP experiments. By directly training not only on the PDB but also on larger sets of protein pairs where direct interactions are confidently known to occur (and not occur), as was done by Motmaen et al. 2023 for peptide-MHC complexes (*79*), it should be possible to increase prediction accuracy across a broad spectrum of interaction modalities.

## Methods

We have built upon our previously developed multi-step bioinformatics and DL pipelines for identifying pairs of interacting proteins within the proteome of an organism (*7, 9*) to improve the scalability and accuracy of predictions. The architecture of the new RF2-Lite, which was trained both on monomeric proteins and protein complexes, is outlined in **Fig. 1A**, and the major steps of the bioinformatics pipeline are listed in **Fig. 1C**.

## Acknowledgments

We thank N.V. Grishin, E. Horvitz, and H. Park for helpful discussions, L. Goldschmidt and A. Guillory for computing resource management, and L. Stewart and L. Stuart for logistical support. Additionally, we are grateful to T.G. Bernhardt, Y.O. Elshenawi, X. Liu, G.V. Mukamolova, K.M. Ottemann, M.L. Reniere, and N.R. Salama for their correspondence and biological expertise.

We acknowledge funding from Bill and Melinda Gates Foundation #OPP1156262 (I.R.H.) and Washington Research Foundation and Translational Research Fund (to M.B.). This work was supported by the National Research Foundation of Korea (NRF) grant funded by the Korea government (MSIT) (No. RS-2023-00210147 to M.B.), and J.Z. was supported by CPRIT training grant RP210041. The Defense Threat Reduction Agency grant HDTRA1-21-1-0007 (A.K. and I.A.), Audacious Project at the Institute for Protein Design (A.K.), Spark Therapeutics (I.A.), and National Institute of Allergy and Infectious Diseases (NIAID) Federal Contracts HHSN272201700059C & 75N93022C00036 (I.A.). NIH R01AI145954 (to J.D.M), Defense Advanced Research Projects Agency Biological Technologies Office Program: Harnessing Enzymatic Activity for Lifesaving Remedies (HEALR) under cooperative agreement No. HR0011-21-2-0012 (to J.D.M), and I-2095-20220331 (to Q.C.) from the Welch Foundation. J.D.M and D.B. are Howard Hughes Medical Institute investigators and Q.C. is a Southwestern Medical Foundation endowed scholar.

## Author contributions

QC and DB conceived the research; IRH and JZ prepared the sequence alignments used in the screen; MB designed and trained RoseTTAFold2-Light; IRH, JZ, JP, IA, and QC designed the PPI screening procedure; IRH and JZ carried out the screen; IRH, JZ, MB, AK, and QC analyzed and presented computational results; YW, CAT, and BAJ conducted laboratory experiments; YW analyzed and presented experimental results; IRH, YW, TW, DTH, SBP, and JDM provided biological insights on specific examples; IRH, YW, SBP, JM, QC, and DB drafted the manuscript; all authors discussed the results and commented on the manuscript

The authors declare they have no competing interests.

