## Supplemental Information for "Essential and virulence-related protein interactions of pathogens revealed through deep learning"

### Table of Contents

**Supplemental Methods and Results** **3**
Multiple sequence alignment generation 3
Genomic database generation 3
Orthologue identification by reciprocal best hits 3
Table S1: Proteome strains and Uniprot accessions 3
Figure S1: Flowchart of reciprocal best hits filtering criterion for complete proteomes 4
Aligning orthologues 4
Figure S2: Representative distribution of monomeric MSA depth 5
Paired multiple sequence alignment generation 5
Figure S3: Paired multiple sequence alignment schematic 6
Proteome filtering 6
Figure S4: Distribution of pMSA depth 7
Figure S5: Distribution of AlphaFold monomer model average pLDDT 7
Identification of virulence factors, essential genes, and uncharacterized proteins 8
Table S2: Statistics for pathogens in our study. 8
Selection and benchmark of the "pilot-set" 9
Positive control selection 9
RoseTTAFold2-Lite 10
Table S3: RoseTTAFold2-Lite training setup 11
Computing contact probability 11
DCA screen implementation 11
RoseTTAFold2-Lite screen implementation 12
AlphaFold screen implementation 12
Benchmark performance of different methods 12
Figure S6: PPI screening methodology performance 13
Figure S7: AlphaFold-multimer distance vs ipTM for PPI identification 14
Interactions between proteins encoded by disparate genes 15
Figure S8: Predicted interactions by genomic distance 15
Figure S9: Predicted unique interactions by genomic distance 16
Figure S10: Normalized fraction of predicted interactions by genomic distance 17
Metadata and Pymol sessions 17
Experimental Methods 18
Plasmid construction 18
Bacterial two-hybrid assay 18
Protein-protein interaction assays with Ni-NTA or VSV-G immunoprecipitation 18
Western blotting 19
Reporting Summary 19
Antibodies 19
Software 19

**Additional Supplemental Tables** 20
Table S4: Recall of filtering pipeline 20
Table S5: Predicted interactions in STRING 20
Table S6: RF2-Lite pilot-set and extended-set pairs by pathogen 21
Table S7: Metadata of experimentally validated interactions by B2H 22
Table S8: Metadata of experimentally validated interactions by Co-IP 23
Table S9: Uniprot annotations of interactions in Figure 3 24
**Additional Supplemental Figures** 25
Figure S11: Experimentally validated pairs, models, and metadata 25
Figure S12: β-galactosidase activity of validated pairs by bacterial-two hybrid 26
Figure S13: PtsH-PtsN negative Co-IP pulldown 27
Figure S14: Uncropped western blot images for Figure 2: I 28
Figure S15: Uncropped western blot images for Figure 2: II 29
Figure S16: Uncropped western blot images for Figure 2: III 30
Figure S17: Uncropped western blot images for Figure 2: IV 31
Figure S18: Uncropped western blot images for Figure 2: V 32
Figure S19: Glucose-6-phosphate 1-dehydrogenase and OPXX cycle protein 33
Figure S20: tRNA 2-thiouridine synthesizing complex (Tus) and MnmA trimers 34
Figure S21: tRNA 2-thiouridine synthesizing complex (Tus) 35
Figure S22: Urease oligomeric assembly generation 36
Figure S23: Urease trimeric interactions 37
Figure S24: Sec translocon orthologous PDB validation and PpiD 38
Figure S25: Sec translocon interactions with CrgA 39
Figure S26: BAM complex orthologous PDB validation 40
Figure S27: BAM complex and SurA interactions 41
Figure S28: BepA putative orthologue identification and Bam/SurA interaction 42
Figure S29: Folding of TolC by BAM complex 43
**References** 44

### Supplemental Methods and Results

#### Multiple sequence alignment generation

##### Genomic database generation

To generate a genomic sequence database, we downloaded 322,271 proteomes and corresponding genomes from NCBI (https://www.ncbi.nlm.nih.gov/genome) and JGI (https://genome.jgi.doe.gov/portal/), available May 2021. We then clustered proteomes by species and selected one to two representatives for each species based on NCBI 'reference' or the largest number of proteins and included all proteomes without species designations. This resulted in a total of 44,871 representative proteomes, which were used to populate the orthologous sequences of the multiple sequence alignments.

##### Orthologue identification by reciprocal best hits

We downloaded the proteomes from uniprot (https://www.uniprot.org/proteomes/) for the 19 bacterial pathogens in this study (table S1). We used these proteins as queries to search for orthologues in the 44,871 representative proteomes/genomes using the reciprocal best hit (rbh) procedure (*1*, *2*) that will be included in the multiple sequence alignment (MSA). Gene duplication events, and fusion events complicate protein evolutionary histories, which may interfere with orthologue identification from sequence similarity to the query alone. Consequently, we use rbh to identify putative orthologues and reduce the likelihood of error. In short, for each query protein in the reference proteome of interest (19 pathogens), we identify the "forward" best hits from each of the representative proteomes, and for each of the "forward" best hits, we identify the "reverse" best hits in the reference proteome of interest by BLAST (*3*) using the criteria shown in fig. S1. For example, given protein *a* in proteome *A*, we identify its "forward" best hits (protein *b*) in proteome *B*, then identify the "reverse" best hits of protein *b* from proteome *B* against proteome *A*. If protein *a* in proteome *A* is among the "reverse" best hits of protein *b*, we designate these proteins *a*,*b* as putative orthologues.

**Table S1: Proteome strains and Uniprot accessions**

| Pathogen (strain) | Uniprot proteome | Pathogen (strain) | Uniprot proteome |
|---|---|---|---|
| **Aca** (PHEA-2) | UP000007477 | **Lpn** (ATCC 33152) | UP000000609 |
| **Bfr** (YCH46) | UP000002197 | **Mge** (G37) | UP000000807 |
| **Bhe** (ATCC 49882) | UP000000421 | **Mtu** (H37Rv) | UP000001584 |
| **Cdi** (630) | UP000001978 | **Nme** (MC58) | UP000000425 |
| **Ctr** (A/HAR-13) | UP000002532 | **Pae** (PA01) | UP000002438 |
| **Eco** (K12) | UP000000625 | **Sau** (PS 47) | UP000008816 |
| **Ftu** (SCHU S4) | UP000001174 | **Spn** (2070335) | UP000002642 |
| **Hpy** (ATCC 700392) | UP000000429 | **Sty** (LT2) | UP000001014 |
| **Lmo** (EGD-e) | UP000000817 | **Vch** (ATCC 39315) | UP000000584 |
| | | **Ype** (CO-92) | UP000000815 |

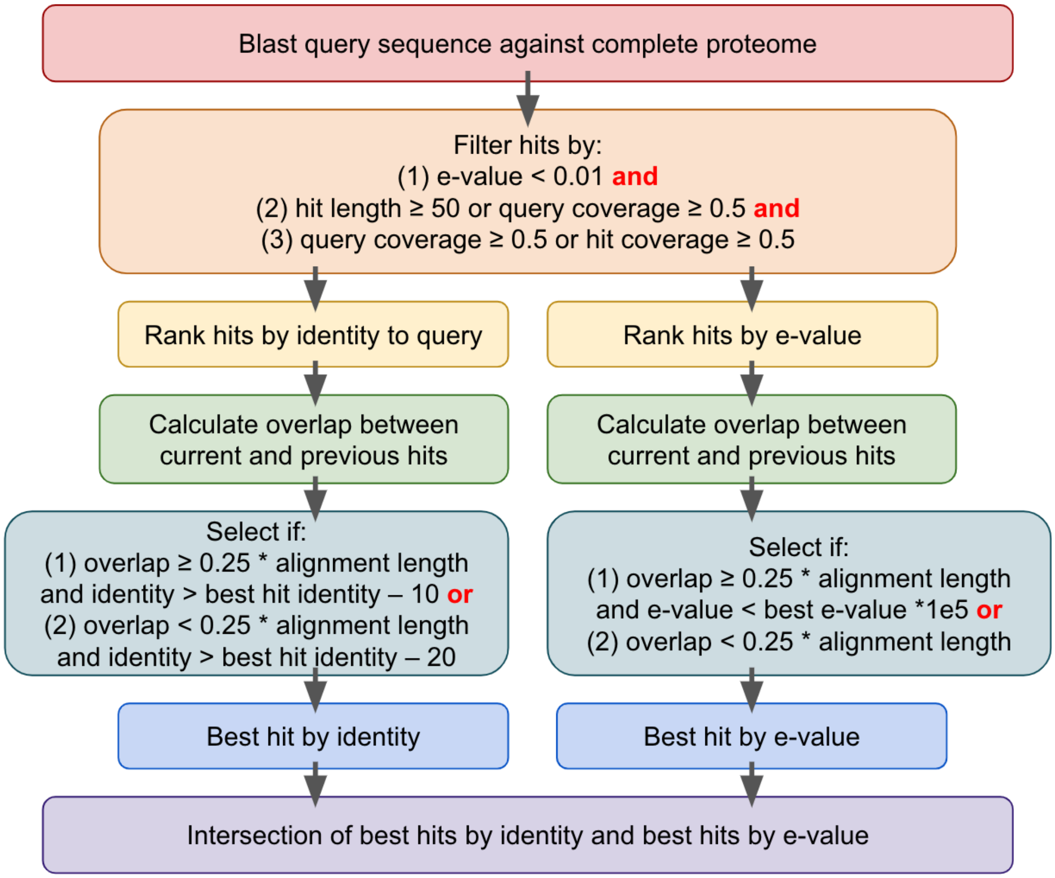


**Figure S1: Flowchart of reciprocal best hits filtering criterion for complete proteomes**

#### Aligning orthologues

Using the above method of orthologue identification from our bacterial protein database, we generate genomic monomeric MSAs for all 57,435 proteins in our dataset (table S2). We applied Hmmer (*4*) to create local sequence alignments of the orthologues identified by rbh. First, we use Phmmer to search each query protein against its orthologues. We filter each MSA to keep highly similar sequences to the query creating a "seed" alignment that was converted into a Hidden Markov Model (HMM) with Hmmbuild and used to align the remaining orthologues with a Hmmsearch (fig. S2). The seed alignment was filtered according to: (a) gap ratio (number of gaps in the alignment / alignment length) < 0.2 and sequence identity (number of identical positions divided by number of aligned positions excluding gaps) ≥ 0.55, (b) gap ratio < 0.35 and sequence identity ≥ 0.4, and (c) gap ratio < 0.5 and sequence identity ≥ 0.25. For each query, the most stringent criteria that selected over 2500 or 25% of all sequences in the orthologous group was used. In the event that Hmmer creates multiple alignments between a query and target, we concatenate the alignments according to their order in the query protein sequence. If aligned segments overlap in the query, we select the overlapping segment from the hit that has a higher sequence identity to the query. We removed positions with gaps in query protein sequences from alignments and sequences with > 50% gap fractions.

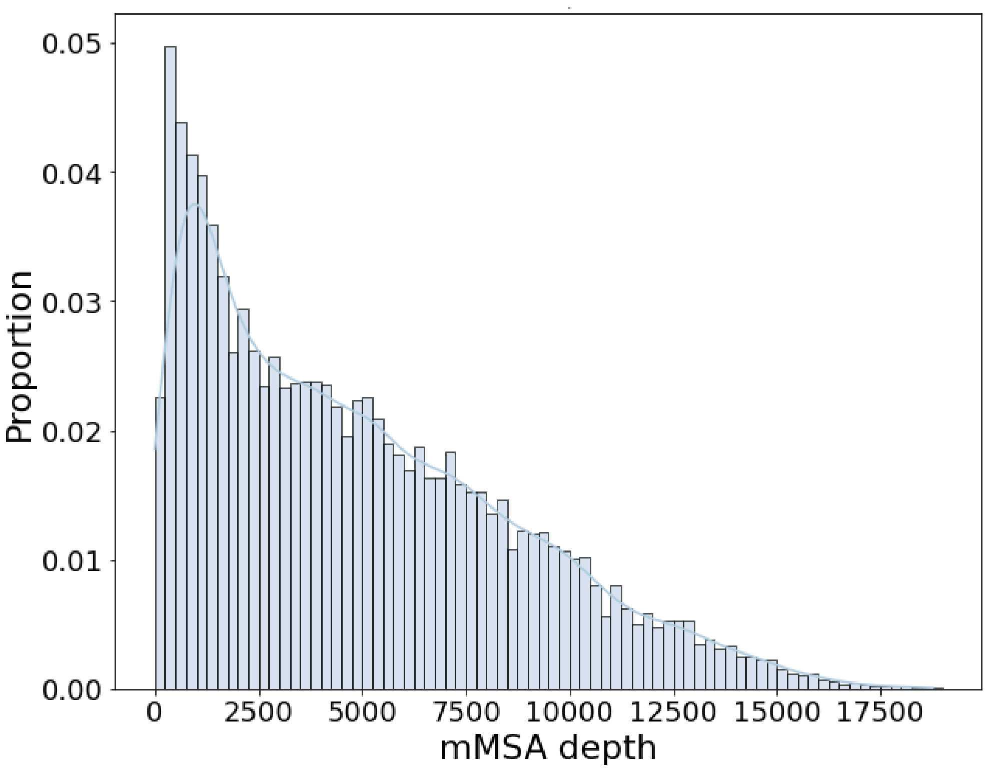


**Figure S2: Representative distribution of monomeric MSA depth**

15,000 proteins across the 19 proteomes were randomly selected to get a representative distribution of monomeric MSA (mMSA) depth. Alignments were filtered with hhfilter at 90% sequence identity and 75% coverage to remove redundancy.

Historically, paralogues have presented a challenge for coevolution-based computational PPI screens; however due to the improvements in performance of RoseTTAFold (RF) and AlphaFold (AF), we find that removal of putative paralogues is no longer necessary to produce high-quality predictions. As such, we did not query a database of paralogues to remove paralogous proteins from our proteome dataset. However, by virtue of rbh criterion, we do not include many potential paralogues in the monomeric multiple sequence alignments.

##### Paired multiple sequence alignment generation

We constructed paired multiple sequence alignments (pMSAs) (fig. S3) for each pair of proteins within each of the 19 bacterial proteomes by concatenating the sequences from the same proteome (each MSA contains a single orthologue per proteome from the bacterial sequence database described above). For each organism, we have $n \times (n - 1) / 2$ pairs of proteins where $n$ is the number of proteins in the proteome. However, with 19 proteomes, the total number of protein pairs was 140,171,142 so we applied several quality control filters as described in subsequent sections along with subsetting out several datasets of interest (i.e., "pilot-set" and "extended-set") to create a computationally tractable study.

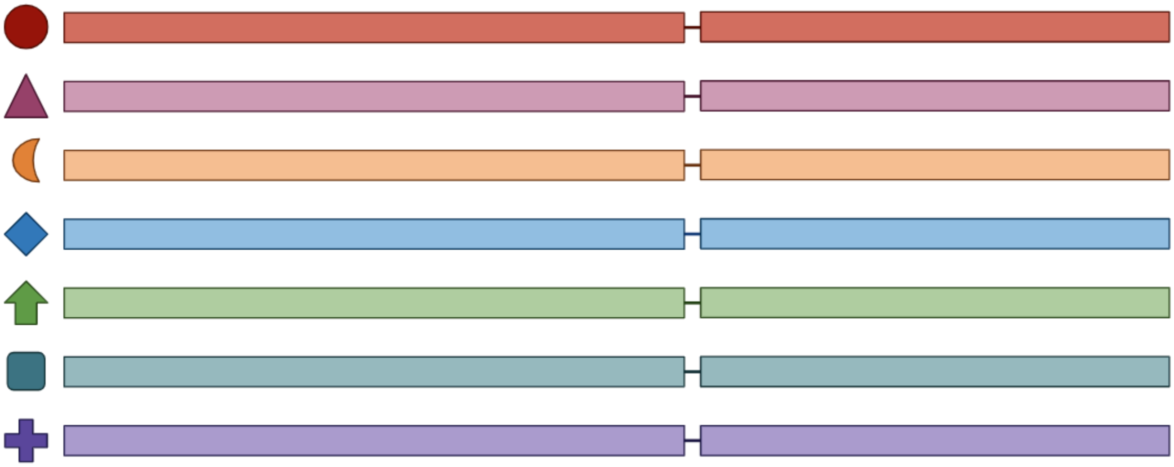

**Figure S3: Paired multiple sequence alignment schematic**

Proteins from the same proteome are colored in the same color. The dash the separation of two proteins and residue indexing gap that is implemented in RoseTTAFold and AlphaFold. However, there is no explicit gap implemented in the pMSAs between two proteins.

#### Proteome filtering

At the core of this computational protein interactome screen lie the evolutionary features extracted from pMSAs and as such, filtering possible candidate proteins or pairs of proteins was necessary as pairs of proteins with shallow multiple sequence alignments are poor candidates for this screen. Additionally, exponentially scaling computational cost with respect to amino acid length limits our ability to predict excessively large proteins and were systematically removed.

We remove pairs with a pMSA depth under 200 sequences after redundancy filtering with hhfilter (*5*) at 75% coverage and 90% sequence identity from our dataset (fig. S4, vertical dashed red line). This resulted in a total of 675,749 pairs excluded across all organisms or approximately 13% of the screened pairs which would otherwise have been added in the "extended-set" pairs. To probe if the proteins in this study could be confidently modeled by AF, we sought to investigate the predicted local distance difference test (pLDDT) of monomeric protein models in our 19 bacterial pathogens. We observe that proteins predicted with low pLDDT or large disordered regions are poor candidates for a combined coevolution and structure-based protein-protein interaction screen. Therefore, we downloaded AlphaFold database (AFDB) V4 (https://alphafold.ebi.ac.uk/) predictions (Nov, 2022) for all proteins that we constructed proteome-wide protein MSAs for. We found that a small number (approximately ~4.4%) of proteins were missing a predicted structure on AFDB. For those that were missing, we computed structures using a local installation of AF if they overlapped with our initial PPI dataset (941 proteins with a length of ≥ 30aa and ≤ 2,000aa). We recorded the average pLDDT from model three with three recycles using our genomic monomeric MSAs (hhfilter coverage 75%, identity 95%).

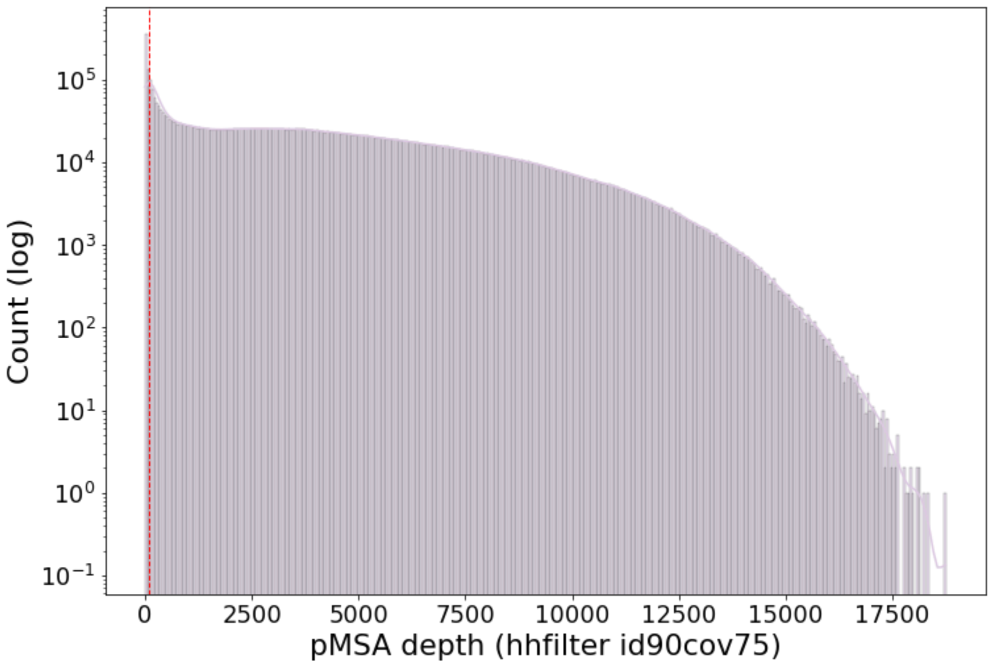


**Figure S4: Distribution of pMSA depth**

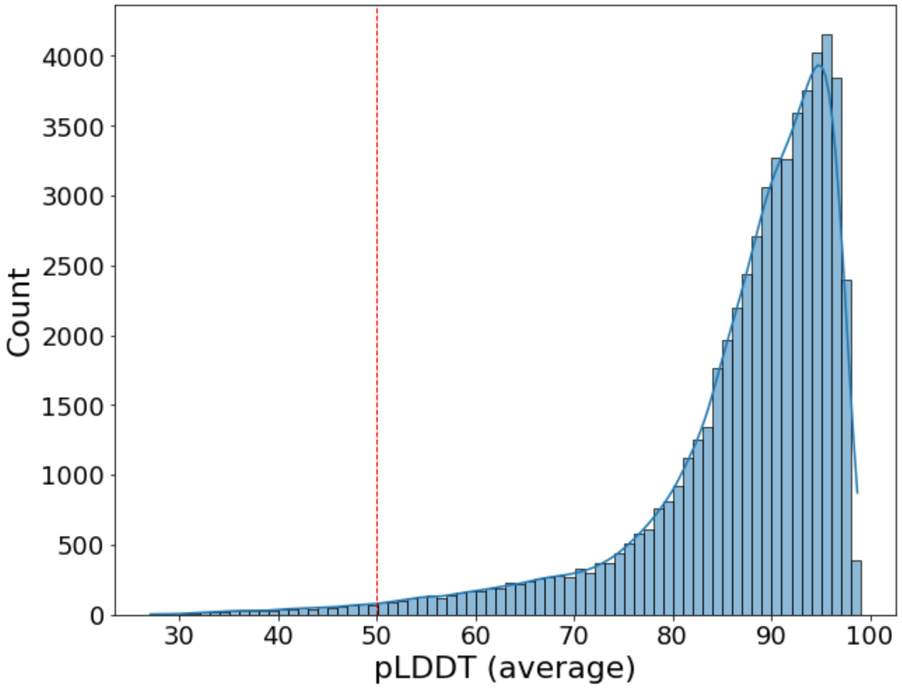


**Figure S5: Distribution of AlphaFold monomer model average pLDDT**

Based on the average pLDDT (fig. S5), we exclude proteins under 50 pLDDT (vertical dashed red line) from our final screen. This resulted in a total of 787 proteins being excluded across all 19 proteomes, or ~1.4% of the total proteins.

#### Identification of virulence factors, essential genes, and uncharacterized proteins

We annotated proteins as essential genes (EG) or virulence factors (VF) based on homology searches to databases. We downloaded the latest release (September 2020) of the DEG bacteria database (*6*). For VFs, we used the 'core dataset' from VFDB (*7*) downloaded in February 2022, which includes genes associated with experimentally verified VFs exclusively in hopes of being more stringent in our VF definition, and Uniprot key words (virulence). Furthermore, any protein that maps to both an EG and VF is designated only EG. Mapping of proteins to the VF and EGs databases was conducted via BLAST sequence similarity searches with a cutoff of at least 90% sequence identity and 80% query and target sequence coverage.

**Table S2: Statistics for pathogens in our study.**

| Abbr. | Organism | Total[1] | Essential genes (unique)[2] | Virulence factors (unique)[3] |
|---|---|---|---|---|
| Aca | *Acinetobacter calcoaceticus* | 3598 | 566 (31) | 34 (6) |
| Bfr | *Bacteroides fragilis* | 4597 | 463 (129) | 0 |
| Bhe | *Bartonella henselae* | 1466 | 3 (0) | 45 (22) |
| Cdi | *Clostridioides difficile* | 3762 | 1 (0) | 13 (10) |
| Ctr | *Chlamydia trachomatis* | 917 | 0 | 113 (95) |
| Eco | *Escherichia coli* | 4262 | 1155 (98) | 51 (15) |
| Ftu | *Francisella tularensis* | 1528 | 495 (46) | 89 (34) |
| Hpy | *Helicobacter pylori* | 1553 | 324 (118) | 113 (47) |
| Lmo | *Listeria monocytogenes* | 2844 | 7 (0) | 35 (14) |
| Lpn | *Legionella pneumophila* | 2930 | 0 | 463 (324) |
| Mge | *Mycoplasma genitalium* | 483 | 381 (101) | 0 |
| Mtu | *Mycobacterium tuberculosis* | 3993 | 1138 (383) | 97 (55) |
| Nme | *Neisseria meningitidis* | 2001 | 534 (43) | 56 (18) |
| Pae | *Pseudomonas aeruginosa* | 5564 | 768 (105) | 289 (88) |
| Sau | *Staphylococcus aureus* | 2889 | 546 (32) | 63 (36) |
| Spn | *Streptococcus pneumoniae* | 2823 | 379 (83) | 32 (21) |
| Sty | *Salmonella typhimurium* | 4533 | 842 (50) | 139 (69) |
| Vch | *Vibrio cholerae* | 3783 | 838 (321) | 141 (43) |
| Ype | *Yersinia pestis* | 3909 | 134 (0) | 120 (22) |
| Total: | | 57,435 | 8574 (1540) | 1893 (919) |

[1]Total number of proteins in the proteome. [2]Number of proteins mapped to essential genes in DEG. [3]Number of proteins mapped to virulence factors in VFDB. In parentheses are the number of unique EG or VF in each organism that lacks orthologues from other pathogens in our study.

Uniprot's protein annotations were updated in 2022 to include protNLM predictions. Consequently, a large number of proteins in uniprot which were previously uncharacterized, or annotated based on homology now have predicted functions. To provide a more robust definition of uncharacterized proteins, we chose to integrate Pfam functional domain mappings and Uniprot's annotations. We downloaded Uniprot Pfam domain mappings for each protein in our dataset. We consider a protein's function to be unknown if all Pfam domain mappings to a given protein were domains of unknown function ("DUF"), "uncharacterized", "unknown", or "putative." Additionally, we manually curated proteins that are annotated as "uncharacterized" in Uniprot based on Pfam domain mappings.

#### Selection and benchmark of the "pilot-set"

All pairs of proteins that include at least one VF or two EG and passed the DCA criterion were included in the "pilot-set." This dataset contains some redundancy with orthologues of VFs or EGs, so for precision/recall benchmarking purposes, we further refined this pilot set to a non-redundant, representative "pilot set" which consists of a single pair per orthologous group selected based on the highest DCA score.

To compare the number of likely novel putative interactions identified in this computational screen, we benchmarked the number of known interaction pairs using our "pilot-set". We identified which of these pairs were known to be interacting based on STRING or contained a structural template based on experimentally solved complexes deposited in the PDB (Fig. 1E). Any pair of proteins from the "pilot-set" that contained a combined string score ≥ 900 and experimental score ≥ 400 were considered to be known interactions by STRING because these pairs are not highly predicted to interact since they are pre-RF/AF screening we included an experimental score cutoff. When mapping to the PDB, we used our "orthologue" mapping criteria as in the metadata section. "Orthologue" structural templates were identified if both proteins aligned to two PDB chains of a deposited complex that were identified to be within 8Å with an interface of at least 5 residues using BLAST with (a) BLAST e-value ≤ 0.00001, and (b) sequence identity ≥ 50% and (c) query or target coverage > 0.5.

#### Positive control selection

The positive control set used in precision/recall calculations was obtained from STRING-db (*8*). We downloaded STRING-db version11 interactions for all 19 pathogens in our dataset. STRING IDs were mapped to uniprot IDs by BLAST sequence identity ≥ 95% and coverage (query and target) ≥ 90%. These STRING interactions were further filtered by an experimental score ≥ 600 and combination score ≥ 990 to reduce the chance of false-positives. The ratio between the positive and negative controls was based on the assumption that each protein, on average, possesses 5 interacting partners out of all other proteins from a species. The negative controls are random pairs with no evidence supporting interaction in STRING. We note that many STRING interactors with experimental support arise from AP/MS experiments which result in possible indirect interactions. Briefly, an indirect interaction may look as such, given proteins A, B, and C, A-B and B-C may be true binary interactions while A-C must be mediated through an interaction with protein B; however, A-C is also denoted as interacting. Since our screen is designed to identify binary interactions, we removed interactions between components of two biological systems: NADH-quinone oxidoreductase and the ribosome based on Uniprot annotations. By removing these large systems, we were able to reduce the number of indirect interactions in our positive control set and more accurately access the power of our screen. Applying a filter specifically for STRING's "physical interactions" is similarly rigorous, with 96% of the pairs selected by our combined and experimental filter passing this additional metric.

#### RoseTTAFold2-Lite

RoseTTAFold2-Lite (RF2-Lite) is an simplified version of the RoseTTAFold2 (*9*) network that has multiple architectural improvements compared to the original RoseTTAFold: (1) use of a three-track architecture with initial coordinates from a template structure, (2) use of biased axial attention to update 2D pair features by considering geometric constraints between residues inferred from the current 3D structure, (3) communication between 1D, 2D, and 3D tracks through attention biasing, and (4) use of recycling that executes the network multiple times with the updated input embeddings based on outputs from the previous cycle. Unlike RoseTTAFold2 that contains 40 3-track blocks, RF2-Lite has 12 3-track blocks to reduce computational costs for large-scale screens.

RF2-Lite was trained based on a mixture of datasets including (1) monomer/homo-oligomer structures in the PDB, (2) hetero-oligomer structures in the PDB (released before August 2nd, 2021), (3) UniRef50 AlphaFold2 structural models having pLDDT > 0.7, and (4) negative protein-protein interaction examples generated by random pairing. The training examples were sampled from each database with a ratio of 1:2:3:2. The model was trained using the masked language model (MLM) loss, distogram (dist) prediction loss, FAPE loss, accuracy estimation loss, bond geometry loss and van der Waals (vdW) energy loss. For the initial round of training, only the first four loss terms were used with crop size 256. After 200 epochs of initial round training, we performed fine-tuning with all the loss terms with crop size 384 for 100 epochs. For the negative interaction examples, we ignored the inter-chain region for FAPE loss calculation and made the network to predict "non-interacting bin" for the inter-chain region for distogram prediction. The training details are summarized in table S3.

The negative training dataset was derived using PDB chains of the same organism from different PDB entries. We removed any protein pairs whose homologs are functionally related according to the STRING database. We sampled a fixed subset of the negative dataset of about the same size as the positive dataset after removing redundancy to provide equivalent data. The total size of both the positive and negative datasets are much larger than the number of training cases in each epoch, where a random sample of positive and negative datasets was used. In our initial model training explorations, we found that incorporating negative pairs alongside positive pairs significantly aids the model in distinguishing true PPIs from random pairings. We maintained an equal ratio of positive to negative pairs to avoid bias that could lead the model to either over-predict or under-predict PPIs. To increase the diversity of protein sequences in the training dataset, we added an equal ratio of monomeric proteins, including protein chains from PDB and high-quality AF models of Uniref representatives. We used a ratio of 1:3 between PDB chains and AF models because the latter has much higher diversity.

**Table S3: RoseTTAFold2-Lite training setup**

| | Initial training | Fine-tuning |
|---|---|---|
| **Crop size** | 256 | 384 |
| **Batch size** | 32 | 32 |
| **Loss function** | 3.0*$Loss_{MLM}$ + 1.0*$Loss_{dist}$ + 10.0*$Loss_{FAPE}$ + 0.1*$Loss_{accuracy}$ | 3.0*$Loss_{MLM}$ + 1.0*$Loss_{dist}$ + 10.0*$Loss_{FAPE}$ + 0.1*$Loss_{accuracy}$ + 0.1*$Loss_{bond}$ + 0.1*$Loss_{vdW}$ |
| **Learning rate & scheduling** | 0.001<br>Linear warm-up for first 1000 optimization steps, then decay learning rate by 0.95 after every 15000 optimization steps | 0.0005<br>No warm-up. Decay learning rate by 0.95 after every 15000 optimization steps |
| **Examples per epoch** | 22400 | 22400 |
| **Number of epochs** | 200 | 100 |

#### Computing contact probability

We used pairs from STRING as positive controls (see above) and randomly selected negative controls (number negative controls = (number of positive controls ✕ number of proteins in proteome) / 5 ) to rank prediction confidence with precision and recall curves.

##### DCA screen implementation

We use our implementation of direct coupling analysis (DCA) that can be computed using GPUs (*10*). We compute the top inter-protein residue-residue DCA score as a measure of protein-protein interaction likelihood. To these scores, we applied average product correction (APC) to raw DCA scores using the formula to reduce false-positive hubs:

$$s'_{i,j} = s_{i,j} - \left(\sum_{m=1}^{m=L1} s_{i,m}\right) \times \left(\sum_{n=1}^{m=L1} s_{n,j}\right) \div \left(\sum_{m=1}^{m=L1} \sum_{n=1}^{m=L1} s_{m,n}\right)$$

Where *m* and *n* represent any residues in the first protein if residues *i* and *j* both belong to the first protein.

$$s'_{i,j} = s_{i,j} - \left(\sum_{m=1}^{m=L2} s_{i,m}\right) \times \left(\sum_{n=1}^{m=L2} s_{n,j}\right) \div \left(\sum_{m=1}^{m=L2} \sum_{m=1}^{m=L2} s_{m,n}\right)$$

Where *m* and *n* represent any residues in the second protein if residues *i* and *j* both belong to the second protein.

$$s'_{i,j} = s_{i,j} - \left(\sum_{m=1}^{m=L2} s_{i,m}\right) \times \left(\sum_{n=1}^{m=L1} s_{n,j}\right) \div \left(\sum_{m=1}^{m=L2} \sum_{m=1}^{m=L1} s_{m,n}\right)$$

Where *m* is any residue in the first protein and *n* is any residue in the second protein if residue *i* is in the first and residue *j* is in the second protein, respectively. Additionally, we remove putative homologous pairs of proteins that are false positives of DCA by removing any pair that contains high scores along the diagonal. All pairs were subject to DCA prior to RF2-Lite.

##### RoseTTAFold2-Lite screen implementation

For both RoseTTAFold2-Lite and AlphaFold, we compute an interaction score ($P_{int}$) based on the summed < 12Å $C_\beta$-$C_\beta$ distance bins between residues *i* and *j* as we used previously (*10*–*12*). Briefly, for a pair of proteins, the matrix (m) contact probability is of the shape (len1 + len2) × (len1 + len2), where len1 and len2 are the lengths of the first and second proteins, respectively. For protein-protein interaction screening, we extract the submatrix m' = m[: len1][len1:len1+len2] which is used for interactions score $P_{int}$.

We previously found that RoseTTAFold-2track (*10*) has the tendency to artificially inflate the contact probability at the C- and N-terminal residues, likely as a result of low confidence secondary structure predictions at these residues, therefore, we continue to exclude the last ten residues of protein 1 and first ten residues of protein 2 from $P_{int}$ with RF2-Lite. However, one difference between RF-2track and RF2-Lite is that RF2-Lite now implements iterative refinement recycling through a network similar to AF. Due to the size of this study, we use three recycles and limit RF2-Lite to 1000 sequences from the pMSA at a time to save computational time and memory. Additionally, due to the increased accuracy of these predictions over RF-2track, we no longer implement average product correction on RF scores.

##### AlphaFold screen implementation

All pairs of proteins that passed RF2-Lite screening at a $P_{int}$ score ≥ 0.05 were subject to AF screening which was run using three recycles and up to 1000 sequences. We use AF with a 200 residue indexing gap to indicate to the AF network that there is no chain connection between the two subunits as previously described (*10*). We reduce the computational cost by producing only a single model (model three) as opposed to the five standard models AF can produce.

#### Benchmark performance of different methods

In addition to the methods described in the above section, we also included AlphaFold-multimer (AF-mm) in our benchmark (*13*). We used the improved v2.0 weights (March 2022 version). We modified the prediction script to enable our custom pMSA input, however, no other modifications were made to the model or prediction scripts and we parse the default distogram output for < 12Å $C_\beta$-$C_\beta$ distance probabilities.

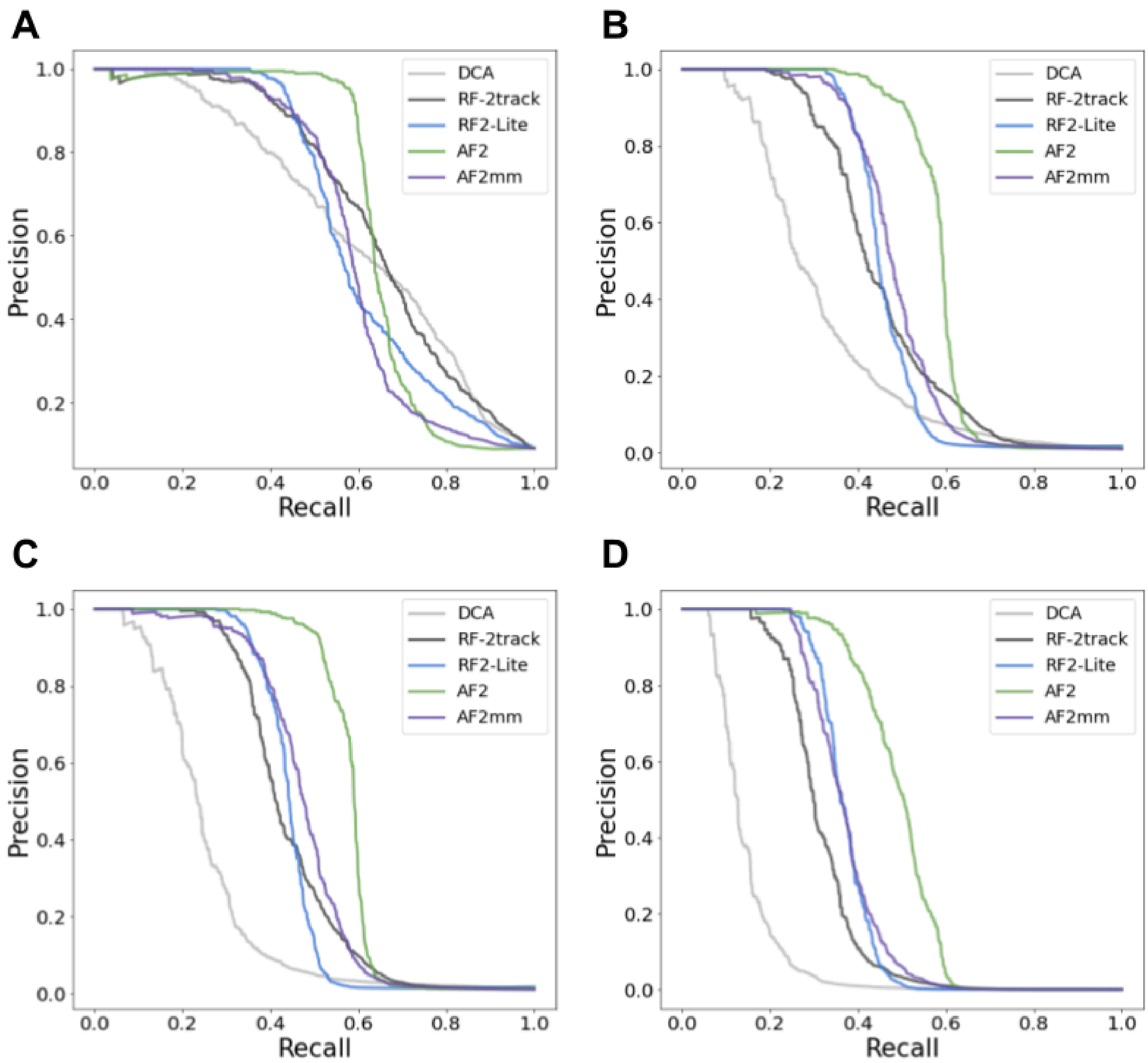


**Figure S6: PPI screening methodology performance**

Precision vs recall of different PPI screening tools. (**A**) Baseline 1000 positive:10,000 negative pair dataset. (**B**) 1000 positives with 100,000 negatives that were generated by randomly sampling 10 data points with 0.1 standard deviation around each of the 10,000 baseline negative examples. (**C**) 1000 positives with 100,000 negatives that were generated by randomly sampling 10 data points with 0.05 standard deviation around each of the 10,000 baseline negative examples. (**D**) 1000 positives with 1,000,000 negatives that were generated by randomly sampling 100 data points with 0.05 standard deviation around each of the 10,000 baseline negative examples.

We tested the performance of several common protein-protein interaction screening methods against RoseTTAFold2-Lite (Fig. 1B; fig. S6). For this analysis, we randomly selected 1000 positive control pairs (excluding NADH-quinone oxidoreductase and ribosomal proteins based on uniprot annotations) from STRING (see above) and 10,000 negative control pairs which were not present in STRING. All pairs are from any of the 19 organisms, contain a monomeric AF pIDDT ≥ 50, pMSA depth ≥ 200 sequences after 75% coverage and 90% identity clustering and range between 100 to 2000 amino acids in length. RF-2 track, RF2-Lite, AF, and AF-mm were provided the same pMSAs, and allowed 1000 sequences, at three recycles where applicable (*note that RF-2 track does not implement recycling*), and we compute statistics on model three. Interaction scores in Fig. 1B and fig. S6 are derived from summing the < 12Å $C_\beta$-$C_\beta$ distance bins as described above. We find that the performance between methods is difficult to distinguish at a ratio of 1:10 true positives to negative examples (fig. S6A), however increasing the number of negative to positive examples to 1:100 (fig. S6B,C) or 1:1,000 (fig. S6D) which are more representative of experimentally identified portions of known interacting vs non-interacting pairs which were found to be around 0.1% (*14*).

The most resource-consuming method in this benchmark is AF2-multimer, which was 4-8 times slower based on our implementation than AF. *Note: we did not fully optimize the code of AF-mm unlike AF, which may contribute to this additional computational burden.* As a result, running AF-mm and AF on 100,000 negative pairs will cost about the same amount of computer resources as detecting the most confident PPIs from 500,000 – 900,000 candidate PPIs (roughly 1-2x the total inference of our PPI screen). Therefore, to reduce computation, we simulated additional negative examples by randomly sampling 10 or 100 data points around each score with a standard deviation of 0.1 or 0.05. We previously compared the RF 2-track against DCA (*10*) with positive and negative control sets (1:1000 ratio). In this work, to conserve resources, we explored alternative strategies to avoid doing extensive negative controls.

Additionally, we compared the performance of < 12Å $C_\beta$-$C_\beta$ distance bins interaction metric to the AF-mm's default 'ranking_confidence' or 'model confidence', which are statistics derived from the predicted TM-score (pTM) and interface pTM (ipTM) scores (fig. S7) with the same benchmarking dataset.

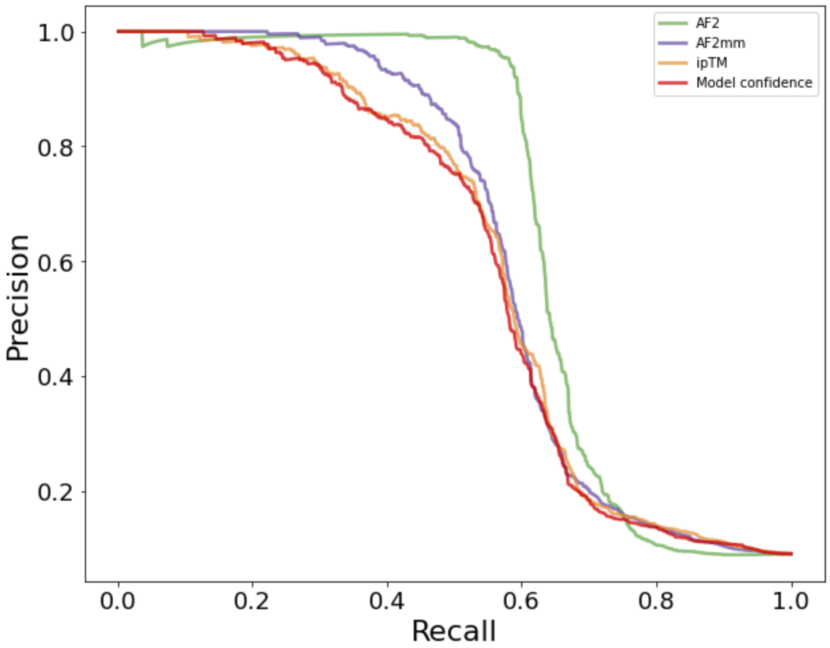


**Figure S7: AlphaFold-multimer distance vs ipTM for PPI identification**

#### Interactions between proteins encoded by disparate genes

To compute the genetic distance between genes on each genome, we downloaded the genome and proteome of each of the 19 bacterial species from NCBI and mapped the uniprot proteomes to these using BLAST. Each uniprot query was assigned to its top respective gene based on e-value ≤ 0.01, sequence identity ≥ 0.8, and query coverage ≥ 0.5. The location within the scaffold was recorded for each gene hit, and the genetic distance between each pair of genes residing on the same scaffold (number of genes between gene1 and gene2) was calculated based on these genome maps accounting for bacterial genomes being circular.

As expected, we observe a large number of neighboring genes score highly by RF2-Lite → AF screening (fig. S8), and those that are highly predicted to interact and are far apart in the genome are less commonly predicted. These interactions that score highly and contain a large genetic distance may be (a) potentially understudied, (b) link between biological pathways, or (c) predicted to interact with a homolog of another protein in the proteome that resides closer to the other protein. To reduce the occurrence of the latter, we remove pairs that have interaction partners with a similar protein < 20 genes from the interacting partner by BLAST e-value ≤ 1e-10. This reduced the number of pairs investigated for genomic distance from 4386 to 2608 pairs that contained an AF score ≥ 0.80 and 2712 down to 1674 pairs at a 0.90 score cutoff (fig. S9).

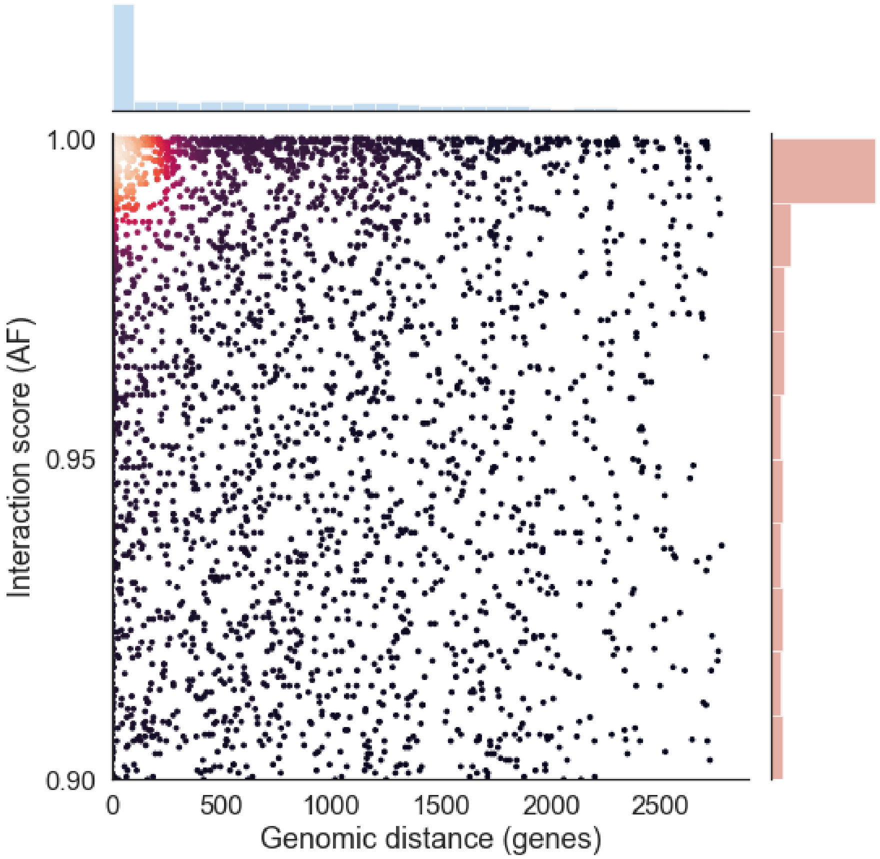


**Figure S8: Predicted interactions by genomic distance**

The primary plot contains the PPI interaction score from the AF screen by the number of genes between the two proteins. Each point represents a pair of proteins which are colored by density computed by gaussian kernel density estimation (KDE). On the side axes of the primary plot, are histograms of these data to better depict the density.

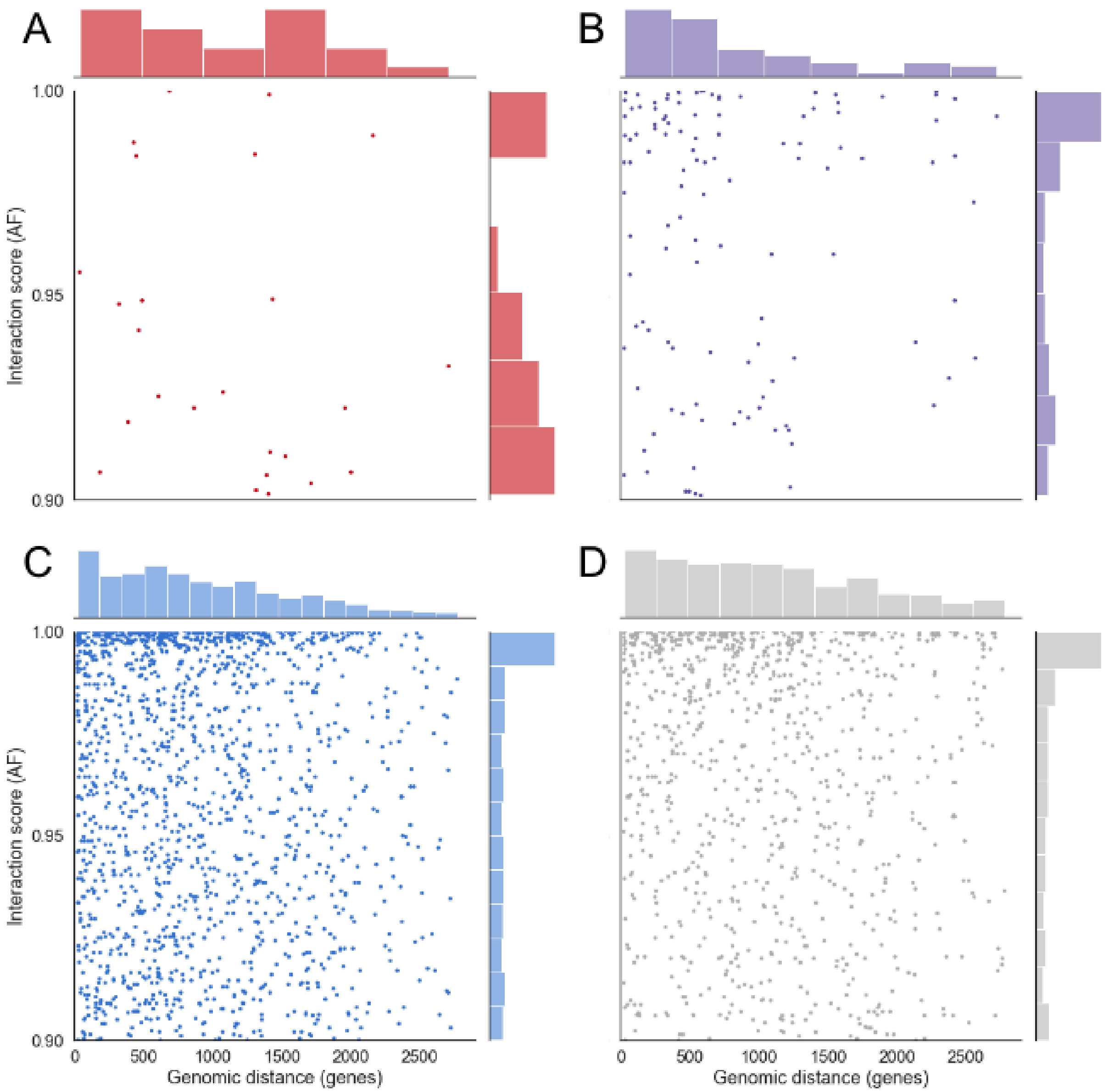


**Figure S9: Predicted unique interactions by genomic distance**

Each primary plot contains the PPI interaction score from the AF screen by the number of genes between the two proteins. Each point represents a pair of proteins. On the side axes of each primary plot, are histograms of these data to better depict the density. (**A**) Interaction between an EG and VF, (**B**) interactions between one VF and non-essential gene or between two VFs, (**C**) interactions between one EG and non-essential gene or between two EGs, and (**D**) interactions between two non-essential, non-virulence factors.

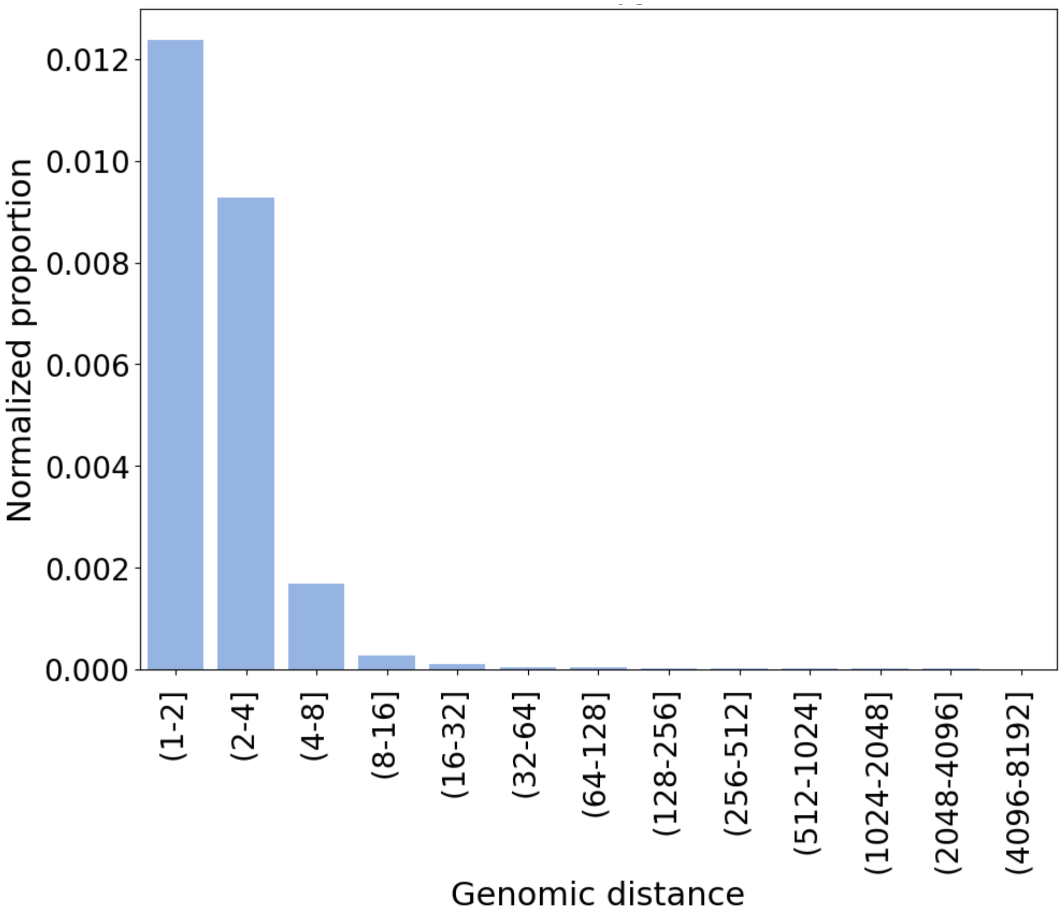


**Figure S10: Normalized fraction of predicted interactions by genomic distance**

Fraction of predicted interactions as a function of genomic distance between protein pairs. Genomic distance is calculated by the minimal number of genes between the coding genes of the two proteins. The proportion of interactions is the number of interactions confidently predicted in our study (AF score > 0.90) normalized to the number of pairs in the 19 pathogen proteomes that fall into each genomic distance bin.

#### Metadata and Pymol sessions

In addition to mapping our protein-protein predictions to STRING-db and genomic location, we also incorporate other auxiliary data into the analysis. We identified the predicted PPIs with experimentally determined structural templates in the PDB. We map our predicted pairs to all pairs of interacting chains (within 12Å with an interface of at least 5 residues) in PDB-deposited complexes using BLAST. The structural templates were split into two categories:

1) Closely ("orthologue") related to a PDB complex if both proteins are aligned to the two interacting PDB chains with BLAST e-value ≤ 0.00001, sequence identity > 0.5, and query coverage > 0.5.
2) Loosely ("homolog") related to a protein complex in the PDB if both proteins can be aligned to interacting chains with BLAST e-value ≤ 0.00001.

To present the modeled protein complexes, we extracted the residue-residue pairs with contact probabilities ≥ 0.6. We render only the top contact probability pair per residue using connecting bars and display the top 10-15 residue contacts for clarity. Furthermore, we remove highly disordered or flexible regions from the rendered structure if they do not participate in the interaction interface. Full structure models containing trimmed regions can be obtained from the ModelArchives.

#### Experimental Methods

##### Plasmid construction

The B2H assay employed the vectors pUT18, pUT18C, pKT25, and pKNT25, which generate N- or C-terminal translational fusions between candidate interaction partners and the T18 and T25 fragments of adenylate cyclase (EUROMEDEX Cat#EUK001, (*15*)). Genes encoding candidate interaction partners were amplified from genomic DNA of the relevant bacterial species using Phusion polymerase and Gibson assembly was used to construct chimeric proteins with T18 or T25 fragments fused to the C-terminus of the protein of interest using pUT18 and pKNT25 vectors. For two interaction pairs, Q9HXJ0-Q9I5H0 and A0A0H2W8E5-A0A384LHF7, chimeric proteins with T18 or T25 fragment fused to the N-terminus of the protein were also generated using pUT18C and pKT25 vectors. Protein expression plasmids used in Ni-NTA pull-down or VSV-G immunoprecipitation were derived from pET-28b(+), and were constructed by amplifying the relevant gene fragments by PCR, and cloning these into the NcoI and XhoI sites of the vector using Gibson assembly. The resulting expression plasmids encode either C-terminally 6xHis-tagged protein or C-terminally VSV-G-tagged protein.

##### Bacterial two-hybrid assay

The bacterial two-hybrid assay was performed essentially as described previously (*16*). *E. coli* BTH101 cells were co-transformed with plasmids encoding the T18 and T25 fragments of *Bordetella pertussis* adenylate cyclase fused to the selected interacting protein pairs. Transformed BTH101 cells were grown in 1 mL LB media supplemented with 0.5 mM IPTG, 150 µg/ml Carbenicillin, and 50 µg/ml Kanamycin in individual wells of a 96-well deep-well plate for ~18 hrs at 30 °C with shaking. The $OD_{600}$ of all cultures was then measured with a Cytation 2 plate reader. 200 µL aliquots of the cultures were transferred to a fresh 96-well plate, and bacterial cells were permeabilized by adding 10 µL chloroform and vortexing vigorously for 10 s. β-Galactosidase activity of 10 µL of the permeabilized cultures was measured using the Galacto-Light Plus™ β-Galactosidase Reporter Gene Assay System (Invitrogen T1011) following the manufacturer's protocol. A luminescence signal indicative of β-galactosidase activity was detected using a Cytation 2 plate reader.

##### Protein-protein interaction assays with Ni-NTA or VSV-G immunoprecipitation

Interactions between predicted PPI pairs were probed using proteins heterologously expressed in *E. coli*. *E .coli* BL21 cells expressing 6xHis or VSV-G-tagged proteins were lysed by sonication in lysis buffer (200 mM NaCl, 50 mM Tris-HCl pH 7.5, 5 mM imidazole, 0.5 mg/mL lysozyme, 25 U/mL benzonase). Cell lysates containing 6xHis or VSV-G-tagged proteins were mixed and the total volume was brought up to 800 mL by equilibration buffer (200 mM NaCl, 50 mM Tris-HCl pH 7.5, 5 mM imidazole). To assess input protein levels, 40 mL of these samples were mixed with Laemmli buffer (Bio-Rad) and boiled for 10 min at 95 ℃, then saved for Western blot analysis. For Ni-NTA pull-down, the remaining protein mixtures were incubated with 50 mL Ni-NTA agarose beads (Qiagen) at 4℃ for 1.5 h with constant rotation. Agarose beads were pelleted by centrifugation at 300 × g for 2 min and washed six times with 1 mL Ni-NTA wash buffer (200 mM NaCl, 50 mM Tris-HCl pH 7.5, and 25 mM imidazole). Proteins bound to the Ni-NTA resin were then eluted by 100 mL Ni-NTA elution buffer (200 mM NaCl, 50 mM Tris-HCl, and 300 mM imidazole). The eluate was mixed with Laemmli loading buffer, boiled and subjected to Western blot analysis.

For VSV-G immunoprecipitation, protein mixtures were incubated with 50 mL anti-VSV-G-agarose beads (Sigma) at 4℃ for 4 h. Agarose beads were pelleted as above and washed six times with 1 mL IP wash buffer (200 mM NaCl, 50 mM Tris-HCl pH 7.5). Proteins bound to the anti-VSV-G-agarose beads were then eluted by 70 mL IP elution buffer (200 mM NaCl, 50 mM Tris-HCl, and 500 µg/mL VSV-G peptides). The eluate was analyzed as above.

##### Western blotting

Input and eluate samples from NTA pull-down and VSV-G immunoprecipitation experiments were separated by SDS-PAGE and transferred to nitrocellulose membranes. Membranes were blocked in TBST (150 mM $NaCl_2$, 10 mM Tris-HCl pH 7.5, and 0.1% v/v Tween-20) with 5% (w/v) bovine serum albumin (BSA) for 1 hr at room temperature. For anti-VSV-G blots, membranes were incubated with anti-VSV-G primary antibody (Sigma) diluted 1:5000 in TBST with 5% (w/v) BSA for 1 hr at room temperature. Blots were then washed 3 times with TBST, followed by incubation with secondary antibody (Goat anti-Rabbit, HRP conjugated, Sigma) diluted 1:5000 in TBST for 1 hr at room temperature. For anti-His blots, membranes were washed twice in TBST after blocking and incubated with anti-His antibody, HRP conjugate (Qiagen) diluted 1:5000 in TBST for 1 hr at room temperature. Finally, blots were washed 3 times with TBST, developed using ECL substrate (BIO-RAD), and visualized using the iBright FL1500 Imaging System (Thermo Fisher).

#### Reporting Summary

##### Antibodies

Anti-VSV-Glycoprotein-Agarose antibody, Mouse monoclonal clone P5D4, Sigma A1970, validated by immunoprecipitation in *E. coli.*

Anti-VSV-G antibody produced in rabbit, Sigma V4888, validated by Western Blot in *E. coli.*

Anti-Rabbit IgG (whole molecule)–Peroxidase antibody produced in goat, Sigma A6154, validated by Western Blot in *E. coli.*

##### Software

Geneious Prime 2023.2.1; Geneious, Software, Newark, New Jersey, USA; Prism 10 for macOS 10.1.1 (270); GraphPad, Software, La Jolla, California, USA; Adobe Illustrator 28.0; Adobe Systems Incorporated, San Jose, California, USA; iBright Analysis Software 5.1.0; Thermo Fisher Scientific Incorporated, Waltham, Massachusetts, USA.

### Additional Supplemental Tables

**Table S4: Recall of filtering pipeline**

| Method(s) of filtering | | | Recall |
|---|---|---|---|
| DCA | | | 4.1% |
| DCA | RF2-Lite | | 28% |
| DCA | RF2-Lite | AF2 | 29% |

Recall of filtering pipeline methods based on 95% precision of pilot-set benchmark based on DCA → RF2-Lite → AF2 successive filtering.

**Table S5: Predicted interactions in STRING**

| Set | STRING (total) | STRING (exp) | New interactions | Existing interactions |
|---|---|---|---|---|
| *Pilot* | 900 | 400 | 381 | 181 |
| *Pilot* | 700 | 400 | 376 | 186 |
| *Pilot* | 400 | 400 | 375 | 187 |
| *Pilot* | 900 | | 287 | 275 |
| *Pilot* | 700 | | 259 | 303 |
| *Pilot* | 400 | | 234 | 328 |
| *All* | 900 | 400 | 2411 | 1202 |
| *All* | 700 | 400 | 2314 | 1299 |
| *All* | 400 | 400 | 2297 | 1316 |
| *All* | 900 | | 1751 | 1862 |
| *All* | 700 | | 1382 | 2231 |
| *All* | 400 | | 1251 | 2362 |

Number of new/known pilot-set and all (extended-set + pilot-set) protein pairs predicted in this screen based on STRING scores at various cutoffs.

Table S6: RF2-Lite pilot-set and extended-set pairs by pathogen

| Abbr. | Organism | Pilot-set | Extended-set | Total |
|---|---|---|---|---|
| Aca | *Acinetobacter calcoaceticus* | 33,094 | 100,173 | 133,267 |
| Bfr | *Bacteroides fragilis* | 13,999 | 55,899 | 69,898 |
| Bhe | *Bartonella henselae* | 344 | 614 | 958 |
| Cdi | *Clostridioides difficile* | 823 | 687 | 1,510 |
| Ctr | *Chlamydia trachomatis* | 1,604 | 2,734 | 4,338 |
| Eco | *Escherichia coli* | 58,417 | 367,586 | 426,003 |
| Ftu | *Francisella tularensis* | 20,327 | 89,404 | 109,73 |
| Hpy | *Helicobacter pylori* | 9,570 | 55,725 | 65,295 |
| Lmo | *Listeria monocytogenes* | 717 | 296,807 | 297,524 |
| Lpn | *Legionella pneumophila* | 7,363 | 50,810 | 58,173 |
| Mge | *Mycoplasma genitalium* | 5,002 | 24,996 | 29,998 |
| Mtu | *Mycobacterium tuberculosis* | 104,177 | 1,096,732 | 1,200,909 |
| Nme | *Neisseria meningitidis* | 20,242 | 96,404 | 116,646 |
| Pae | *Pseudomonas aeruginosa* | 83,052 | 1,235,506 | 1,318,558 |
| Sau | *Staphylococcus aureus* | 25,257 | 57,957 | 83,21 |
| Spn | *Streptococcus pneumoniae* | 18,387 | 45,283 | 63,670 |
| Sty | *Salmonella typhimurium* | 31,144 | 127,260 | 158,40 |
| Vch | *Vibrio cholerae* | 19,698 | 98,932 | 118,630 |
| Ype | *Yersinia pestis* | 4,092 | 20,707 | 24,799 |
| Total: | | 457,310 | 3,824,215 | 4,281,525 |

Pilot-set and extended-set protein pairs screened in this study. All pilot-set were selected from the top scoring pairs by DCA that were annotated as virulence factors or essential genes and run through RF2-Lite.

**Table S7: Metadata of experimentally validated interactions by B2H**

| Org. | Uniprot | Locus | Gene | Annotations |
|---|---|---|---|---|
| Lpn | Q5ZRK0 | lpg2881 | - | Iron-sulfur cluster binding protein |
| | Q5ZYK1 | lpg0371 | - | UPF0125 protein lpg0371 |
| Pae | Q9HX22 | PA4005 | RsfS | Ribosomal silencing factor RsfS |
| | Q9HX38 | PA3981 | YbeZ | PhoH-like protein domain-containing protein |
| Lmo | Q8Y3X6 | lmo2703 | - | Nucleoid-associated protein lmo2703 |
| | Q8Y695 | - | Ffh | Signal recognition particle protein |
| Lmo | P60415 | lmo2054 | - | UPF0298 protein lmo2054 |
| | Q928N1 | lmo2402 | - | Lmo2402 protein |
| Pae | Q9HXJ0 | PA3812 | IscA | Iron-binding protein IscA |
| | Q9I5H0 | PA0759 | YgfZ | Folate-binding protein YgfZ |
| Ype | A0A0H2W8E5 | YPO3902 | - | Putative magnesium chelatase family protein |
| | A0A384LHF7 | YPO0243 | - | DprA winged helix domain-containing protein |
| Pae | Q9HU12 | PA5177 | - | Probable hydrolase |
| | Q9HX75 | PA3941 | - | Uncharacterized protein |
| Sty | P26401 | STM2087 | RfbV | Abequosyltransferase RfbV |
| | Q8ZL53 | STM3707 | YibD | Putative glycosyltransferase |
| Vch | Q9KM04 | VC_A0585 | - | Glutathione S-transferase, putative |
| | Q9KUE5 | VC_0576 | - | Stringent starvation protein A |
| Lpn | Q5ZRT6 | lpg2792 | TpiA | Triosephosphate isomerase |
| | Q5ZTZ8 | lpg2010 | Gmk | Guanylate kinase |
| Lmo | P0A4L3 | lmo1233 | TrxA | Thioredoxin |
| | Q92A74 | - | Hup | Hup protein |

Experimentally validated interaction pairs by B2H displayed in Fig. 2 and figs. S11-12 with organism abbreviations (see above), uniprot ID, gene locus, name, and annotations.

#### Table S8: Metadata of experimentally validated interactions by Co-IP

| Org. | Uniprot | Locus Gene | Score | Contacts | Globularity | Genetic distance | STRING | Annotations |
|---|---|---|---|---|---|---|---|---|
| Eco | P0A887<br>P0AAZ7 | ubiE<br>ycaR | 0.99 | 79 | 98%<br>92% | 1408 | 287 | Ubiquinone/menaquinone biosynthesis C-methyltransferase<br>IspA family inner membrane protein; Involved in cell division |
| Pae | Q9HWS2<br>Q9HWS3 | PA4106<br>PA4105 | 1.0 | 203 | 98%<br>98% | 2 | na | UPF0276 protein PA4106<br>Putative DNA-binding domain-containing protein |
| Pae | P72139<br>Q9HZ78 | hisF2<br>wbpG | 0.99 | 99 | 98%<br>100% | 2 | na | Putative imidazole glycerol phosphate synthase subunit hisF2<br>LPS biosynthesis protein WbpG |
| Lpn | Q5ZRK0<br>Q5ZYK1 | lpg2881<br>lpg0371 | 1.0 | 114 | 86%<br>83% | 475 | 573 | Iron-sulfur cluster binding protein<br>UPF0125 protein lpg0371 |
| Pae | Q9HVV2<br>Q9HVV4 | ptsH<br>ptsN | 0.96 | 54 | 100%<br>100% | 3 | 959 | Phosphocarrier protein HPr<br>Nitrogen regulatory protein |
| Lpn | Q5ZW63<br>Q5ZX88 | FlgJ<br>Ttg2D | 0.94 | 54 | 87%<br>90% | 375 | na | Muramidase, peptidoglycan hydrolase FlgJ<br>Signal peptide protein, toluene tolerance protein Ttg2D |

Experimentally validated interaction pairs by Co-IP displayed in Fig. 2 with organism abbreviations (see above), uniprot ID, locus/gene, AFscores, number of contacts at interface, globularity, genetic distance, STRING combined score, and uniprot annotations. The four pairs above the green line were found to interact by Co-IP; the two pairs below the red line were not detected to interact by Co-IP.

#### Table S9: Uniprot annotations of interactions in Figure 3

| Fig. 3 | Org. | Pair | Protein 1 | Protein 2 |
|---|---|---|---|---|
| A | Mtu | OpcA-G6PD2 (zwf2) | OXPP cycle protein OpcA | Glucose-6-phosphate 1-dehydrogenase 2 (G6PD 2) |
| B | Eco | SapD-SapB | Putrescine export system ATP-binding protein SapD | Putrescine export system permease protein SapB |
| C | Eco | MreB-RodZ | Cell shape-determining protein MreB | Cytoskeleton protein RodZ |
| D | Mtu | PonA-MutT4 | Penicillin-binding protein 1A (PBP-1A) | Putative mutator protein MutT4 |
| E | Pae | PA3801-PpiD | Ancillary SecYEG translocon subunit | Peptidyl-prolyl cis-trans isomerase D |
| F | Pae | RpsK-YbeY | 30S ribosomal protein S11 | Endoribonuclease YbeY |
| G | Bfr | RimH-RsmH | Ribosome maturation factor RimM | Ribosomal RNA small subunit methyltransferase H |
| H | Ftu | MnmA-RimP | tRNA-specific 2-thiouridylase MnmA | Ribosome maturation factor RimP |
| I | Eco | TsaE-PaaX | tRNA threonylcarbamoyl- adenosine biosynthesis protein TsaE | Transcriptional repressor PaaX |
| J | Aca | Ttg2D-VacJ | Putative toluene tolerance protein | VacJ family lipoprotein |
| K | Pae | FliA-FlgM | RNA polymerase sigma factor FliA (RNA polymerase sigma factor for flagellar operon) | Negative regulator of flagellin synthesis (Anti-sigma-28 factor) |
| L | Mtu | DrrC-DrrA | Probable doxorubicin resistance ABC transporter permease protein DrrC | Doxorubicin resistance ATP-binding protein DrrA |
| M | Pae | MucC-RnfA | Positive regulator for alginate biosynthesis MucC | Ion-translocating oxidoreductase complex subunit A |
| N | Spn | AMCSP13_001135 AMCSP13_000945 | Fibronectin/fibrinogen binding domain protein | DNA gyrase subunit B domain protein |
| O | Lpn | CcmE-CcmC | Cytochrome c-type biogenesis protein CcmE | Heme exporter protein C |
| P | Pae | FlgM-FliS | Negative regulator of flagellin synthesis (Anti-sigma-28 factor) | Flagellar secretion chaperone FliSB |
| Q | Pae | Pa1955-Pa1952 | Adhesin | Fimbrial biogenesis outer membrane usher protein |
| R | Lpn | FeoA-RsfS | Ferrous iron transporter A | Ribosomal silencing factor RsfS |
| S | Lpn | lpg1546-LptE | Fimbrial biogenesis and twitching motility protein PilF | LPS-assembly lipoprotein LptE |
| T | Pae | RhlA-LolA | 3-(3-hydroxydecanoyloxy) decanoate synthase | Outer-membrane lipoprotein carrier protein |
| U | Pae | ilvC-PA2823 | Ketol-acid reductoisomerase | DUF815 domain-containing protein |
| V | Pae | PA1065-glpE | DUF488 domain-containing protein | Thiosulfate sulfurtransferase GlpE |
| W | Pae | PA4431-PA0388 | Ubiquinol-cytochrome c reductase iron-sulfur subunit | DUF4426 domain-containing protein |
| X | Mtu | Rv0883c-FtsZ | Uncharacterized protein | Cell division protein FtsZ |
| Y | Mtu | RelA-Rv1312 | Bifunctional (p)ppGpp synthase/hydrolase RelA | Uncharacterized protein |

Predicted interaction pairs displayed in Fig. 3 with uniprot mapping of gene name and protein annotations.

#### Additional Supplemental Figures

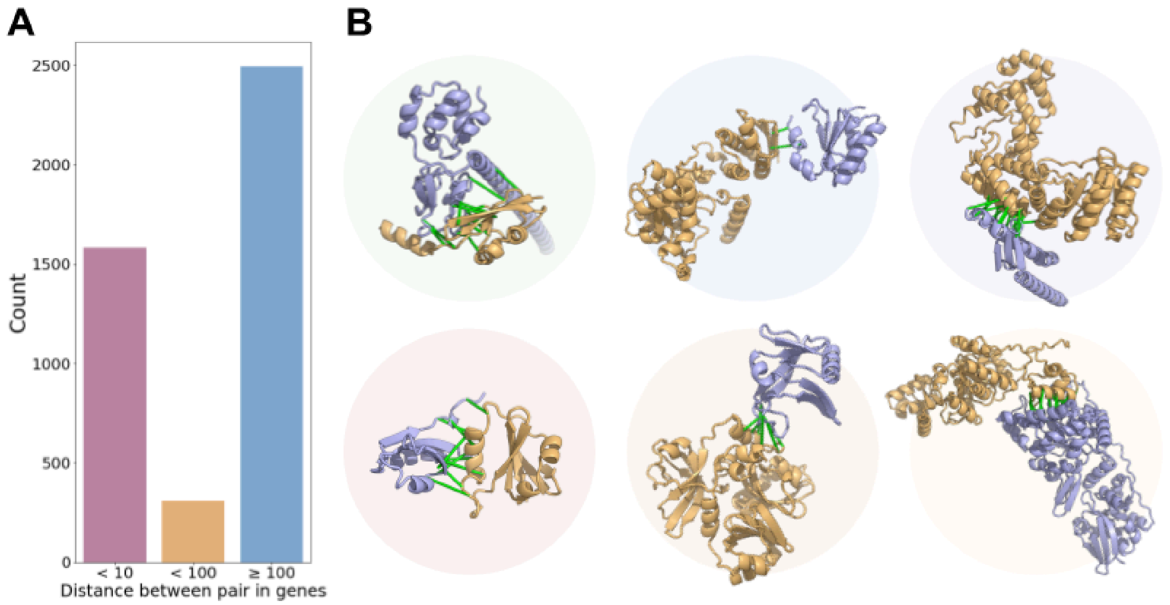


**C**

| | Distance | STRING | DCA | GREMLIN | RF2-Lite | AF | B2H |
|---|---|---|---|---|---|---|---|
| Q5ZRK0_Q5ZYK1 | 475 | 573 | 0.23 | 1 | 1.0 | 1.0 | 0.98 |
| Q9HX22_Q9HX38 | 23 | 736 | 0.77 | 1 | 0.77 | 0.61 | 0.56 |
| Q8Y3X6_Q8Y695 | 899 | 445 | 0.17 | 0.81 | 0.81 | 0.88 | 0.51 |
| P60415_Q928N1 | 347 | 522 | 0.09 | 0.87 | 0.92 | 0.99 | 0.06 |
| Q9HXJ0_Q9I5H0 | 2508 | 445 | 0.55 | 0.98 | 0.79 | 0.81 | 0.03 |
| A0A0H2W8E5_A0A384LHF7 | 419 | 780 | 0.64 | 0.99 | 0.64 | 1.0 | 0.03 |
| Q9HU12_Q9HX75 | 1227 | | 0.14 | 0.83 | 0.01 | 0.27 | 0.03 |
| P26401_Q8ZL53 | 1557 | | 0.12 | 0.74 | 0.005 | 0.07 | 0.11 |
| Q9KM04_Q9KUE5 | | | 0.13 | 0.62 | 0.001 | 0.03 | 0.03 |
| Q5ZRT6_Q5ZTZ8 | 758 | 677 | 0.95 | 1 | 0.003 | 0.06 | 0.04 |
| P0A4L3_Q92A74 | 700 | 295 | 0.38 | 0.74 | 0.04 | 0.03 | 0.07 |

**Figure S11: Experimentally validated pairs, models, and metadata**

Experimentally validated predicted interacting protein pairs encoded by distant genes. (**A**) Distance between predicted pairs of genes ≥ 0.8 AF score (~80% precision). (**B**) Predicted dimeric protein models with AF; left to right: Q5ZRK0-Q5ZYK1Q (green circle), Q9HX22-Q9HX38 (blue circle), Q8Y3X6-Q8Y695 (purple circle), P60415-Q928N1 (red circle), Q9HXJ0-Q9I5H0 (orange circle), A0A0H2W8E5-A0A384LHF7 (yellow circle); green bars between unique predicted interface residues ≤ 12Å. (**C**) Metadata table for pairs containing genetic distance, combined STRING scores, DCA and GREMLIN normalized to precision/recall curve, RF2-Lite and AF scores, and bacterial-two hybrid normalized to positive control (fig. S11). Annotations in table S7.

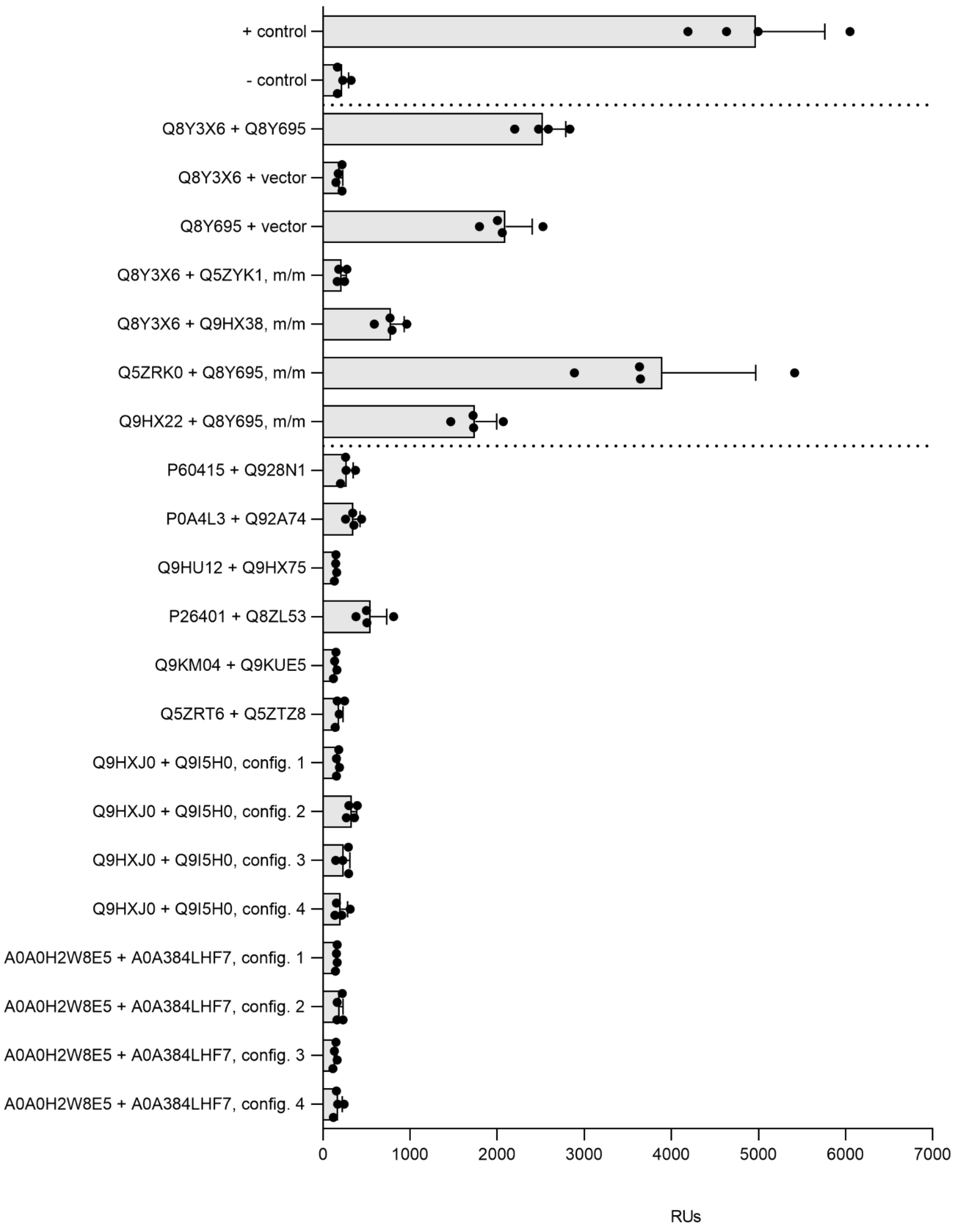


**Figure S12: β-galactosidase activity of validated pairs by bacterial-two hybrid**

Bacterial two-hybrid assays support a lack of interaction between coevolved proteins not predicted to interact by deep learning PPI methods. Assay and controls are as described in Fig. 2A. Config.1 ~ Config.4, different combinations of N-terminal or C-terminal fusions of T18 or T25 fragments with proteins of interest. Error bars indicate ±s.d. (n = 2 biological replicates each with n = 2 technical replicates). Annotations for proteins in table S6.

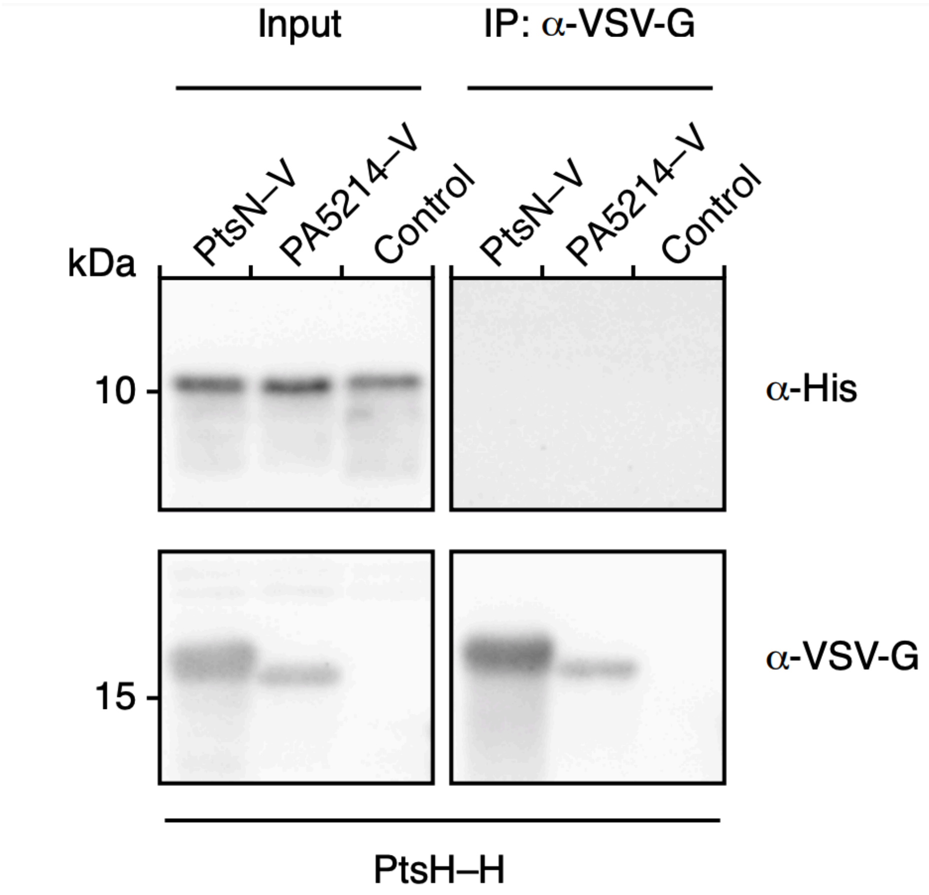


**Figure S13: PtsH-PtsN negative Co-IP pulldown**

α-VSV-G blot for PtsH-PtsN pulldown showing no signal by Co-IP despite strong string score and high PPI score suggesting potential challenges in experimentally validating some putative interaction pairs.

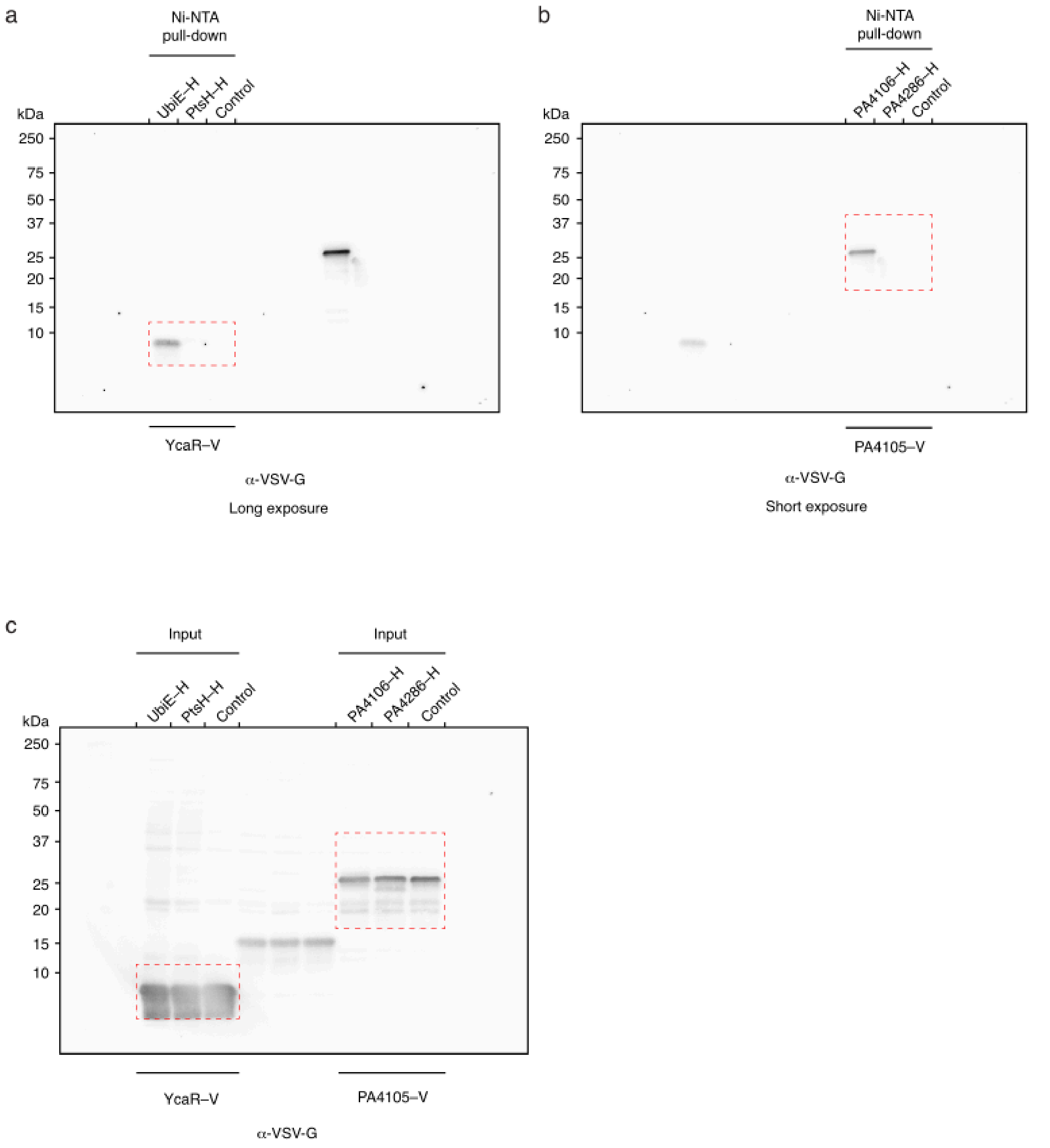


**Figure S14: Uncropped western blot images for Figure 2: I**

α-VSV-G blots for Fig 2a,b. The cropped region of each gel included in the main text figure is denoted by red dash boxes.

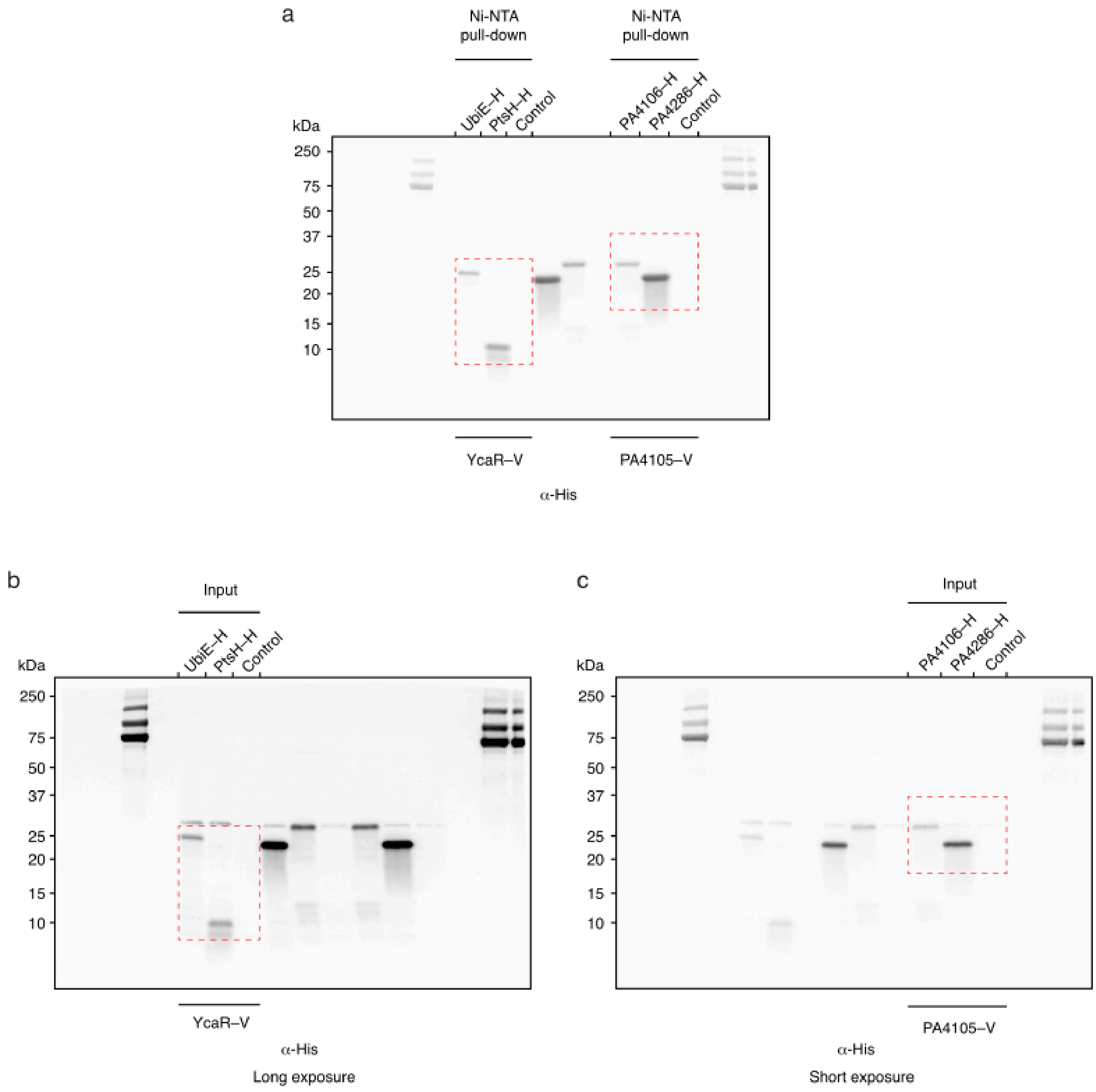


**Figure S15: Uncropped western blot images for Figure 2: II**

α-His-G blots for Fig 2a,b. The cropped region of each gel included in the main text figure is denoted by red dash boxes.

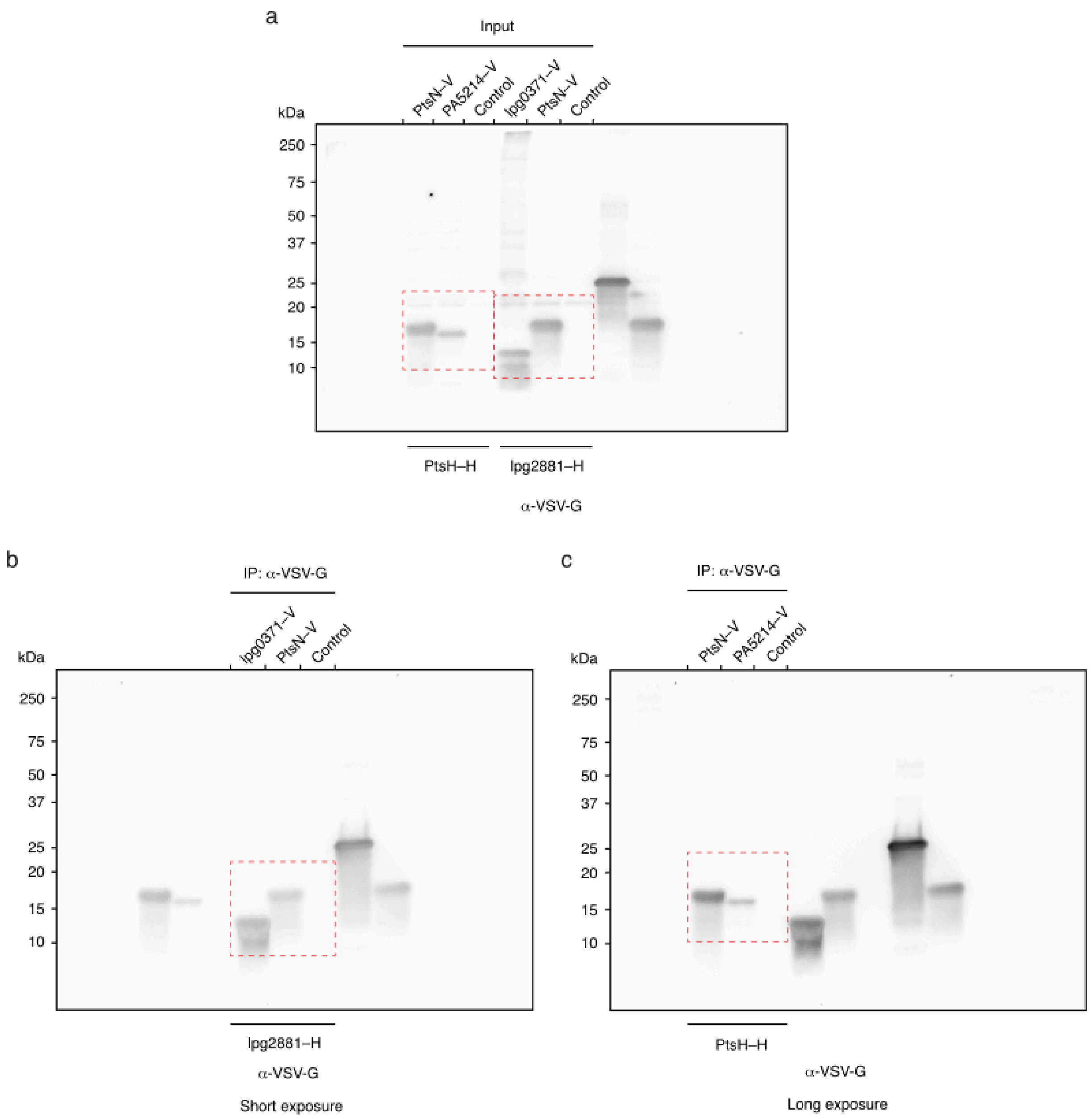


**Figure S16: Uncropped western blot images for Figure 2: III**

α-VSV-G blots for Fig 2d and supplemental figure S13. The cropped region of each gel included in the main text figure is denoted by red dash boxes.

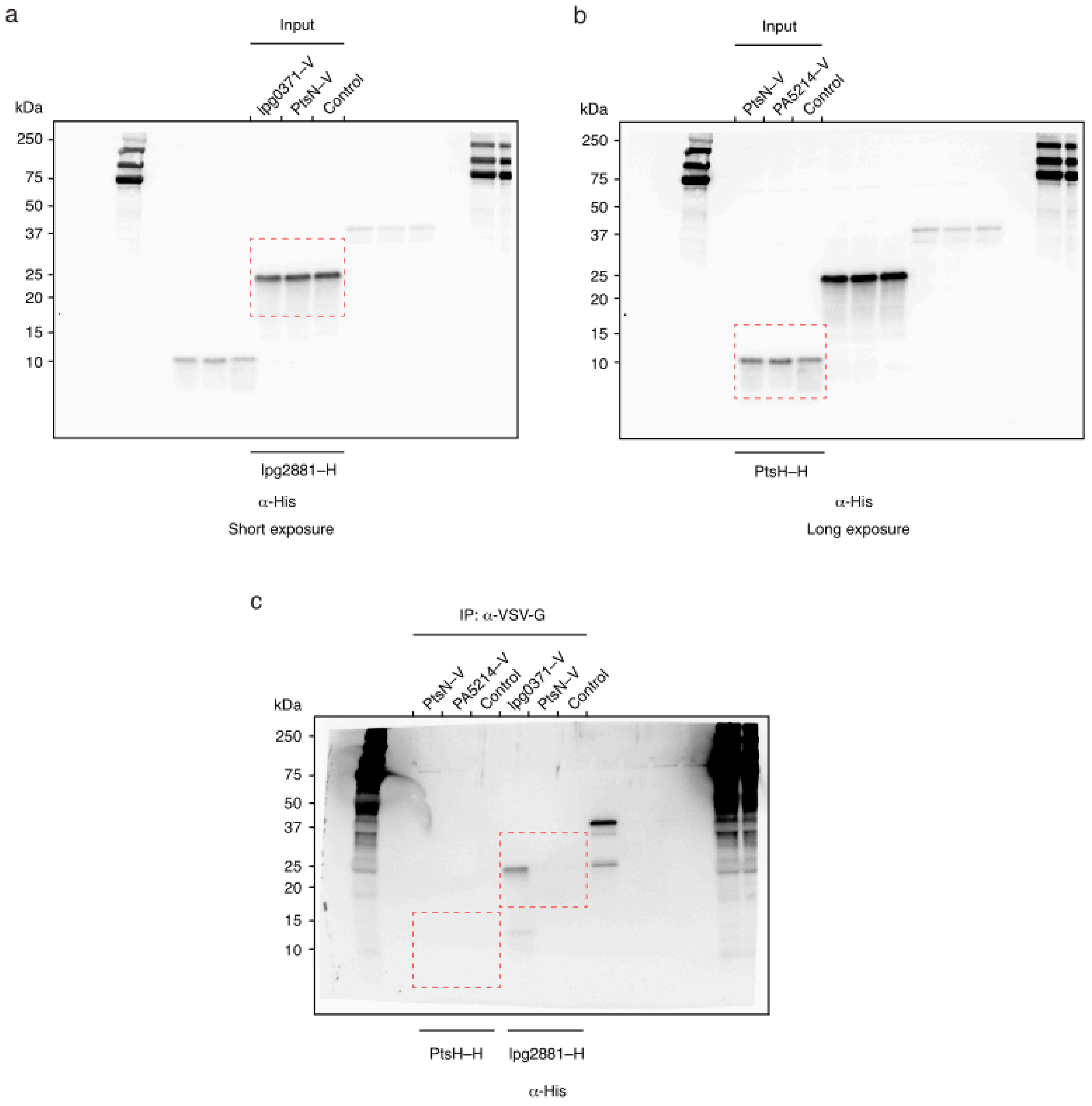


**Figure S17: Uncropped western blot images for Figure 2: IV**

α-His-G blots for Fig 2d and supplemental figure S13. The cropped region of each gel included in the main text figure is denoted by red dash boxes.

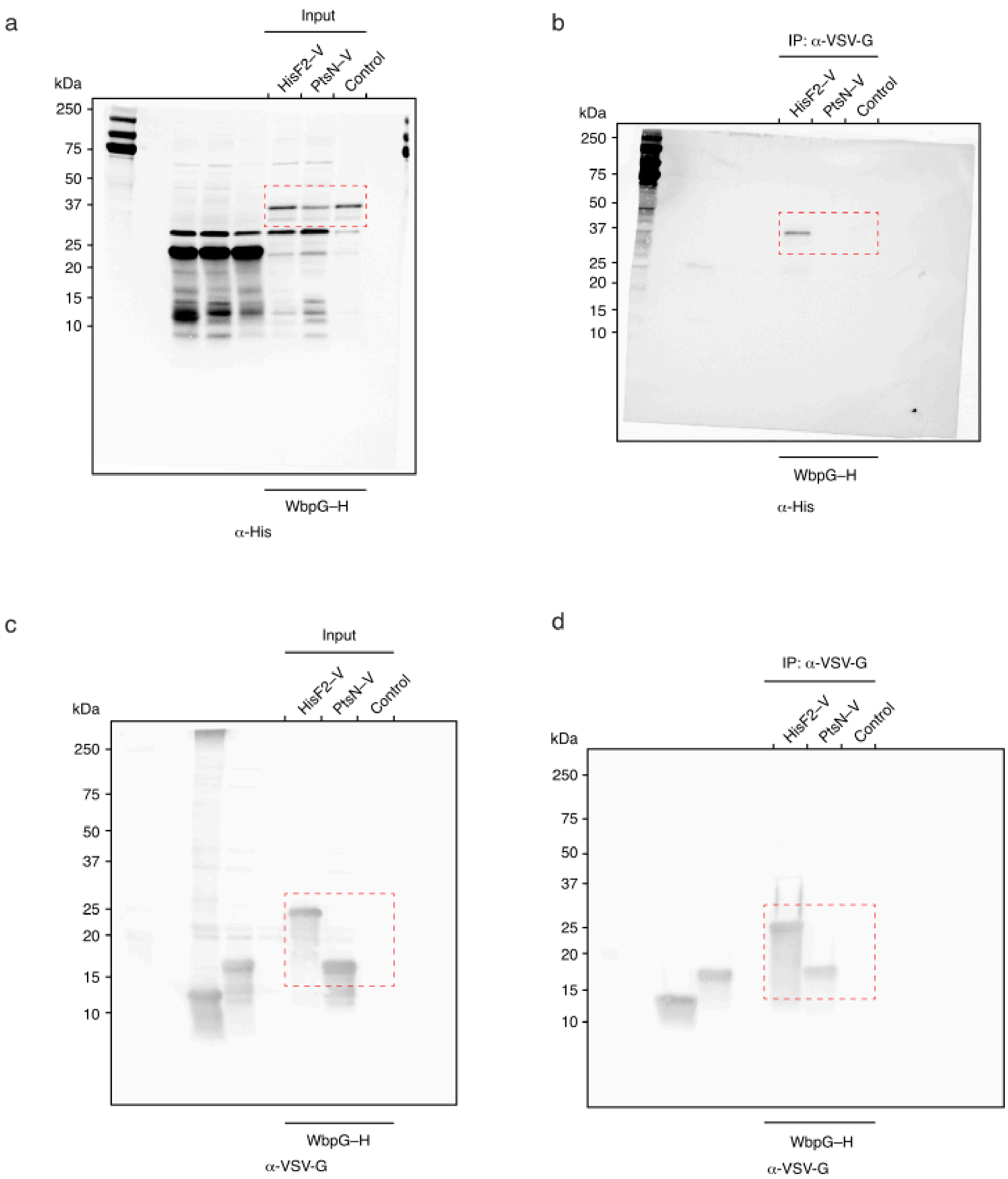


**Figure S18: Uncropped western blot images for Figure 2: V**

α-His and α-VSV-G blots for Fig 2e. The cropped region of each gel included in the main text figure is denoted by red dash boxes.

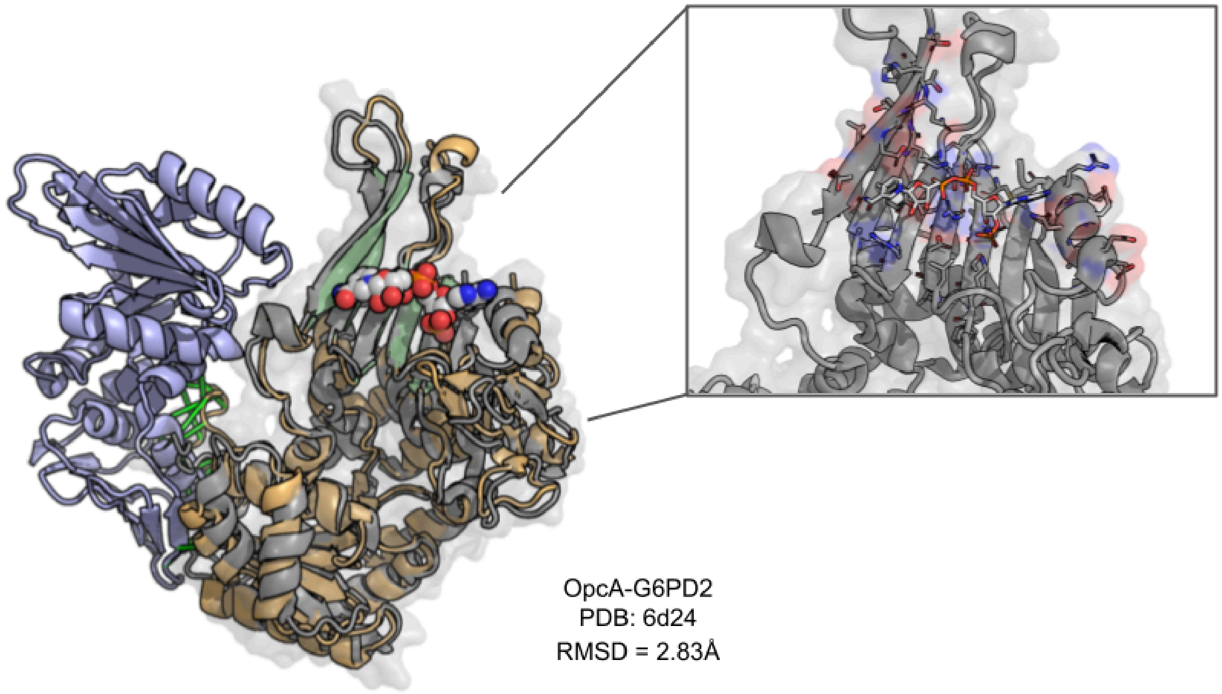


**Figure S19: Glucose-6-phosphate 1-dehydrogenase and OPXX cycle protein**

Predicted interaction of *M. tuberculosis* glucose-6-phosphate 1-dehydrogenase 2 (G6PD2) rendered in gold and OXPP cycle protein OpcA rendered in blue. The active site of the predicted structure is highlighted in green based on PDB annotation. The grey structure is an overlaid structure of G6PD (PDB: 6D24) (*17*), focus box displays the PDB residues and NAD moiety.

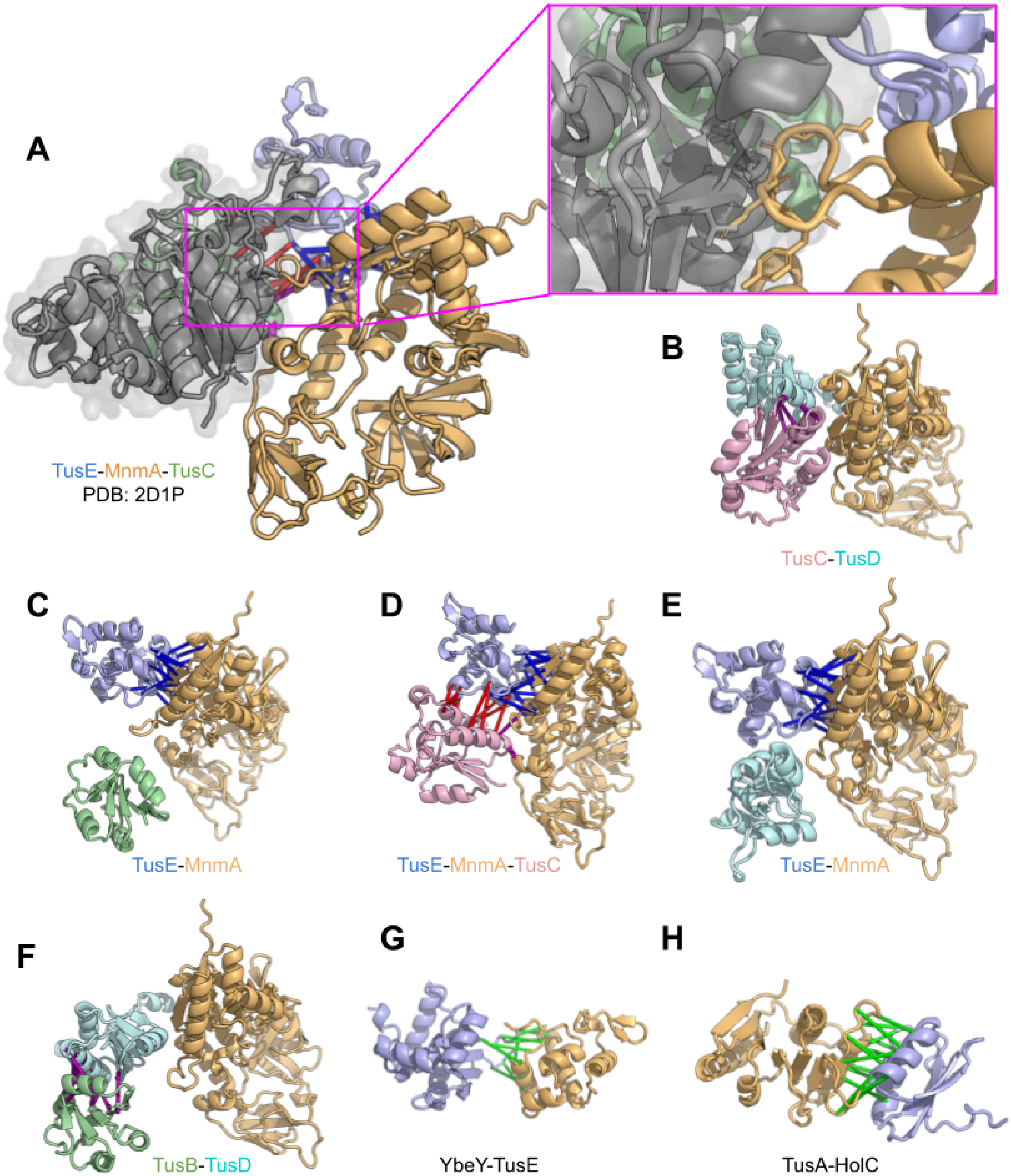


**Figure S20: tRNA 2-thiouridine synthesizing complex (Tus) and MnmA trimers**

Predictions of interactions with Tus in *E. coli*. (A) The modeled trimer of TusE-MnmA-TusC with TusBCD overlay (PDB: 2D1P) (*18*) aligned to TusC showing that while MnmA may interact with TusEC, there are steric clashes with TusBCD; although it should be noted that there may be some flexibility to accommodate this interface which we are unable to capture though overlaid structures. (B-F) Attempted trimeric combinatorial interaction modeling of TusB, TusC, TusD, and TusE with MnmA; below each model the components which were predicted to interact are named. (G) Predicted interaction between YbeY-TusE. (H) Predicted interaction between TusA-HolC.

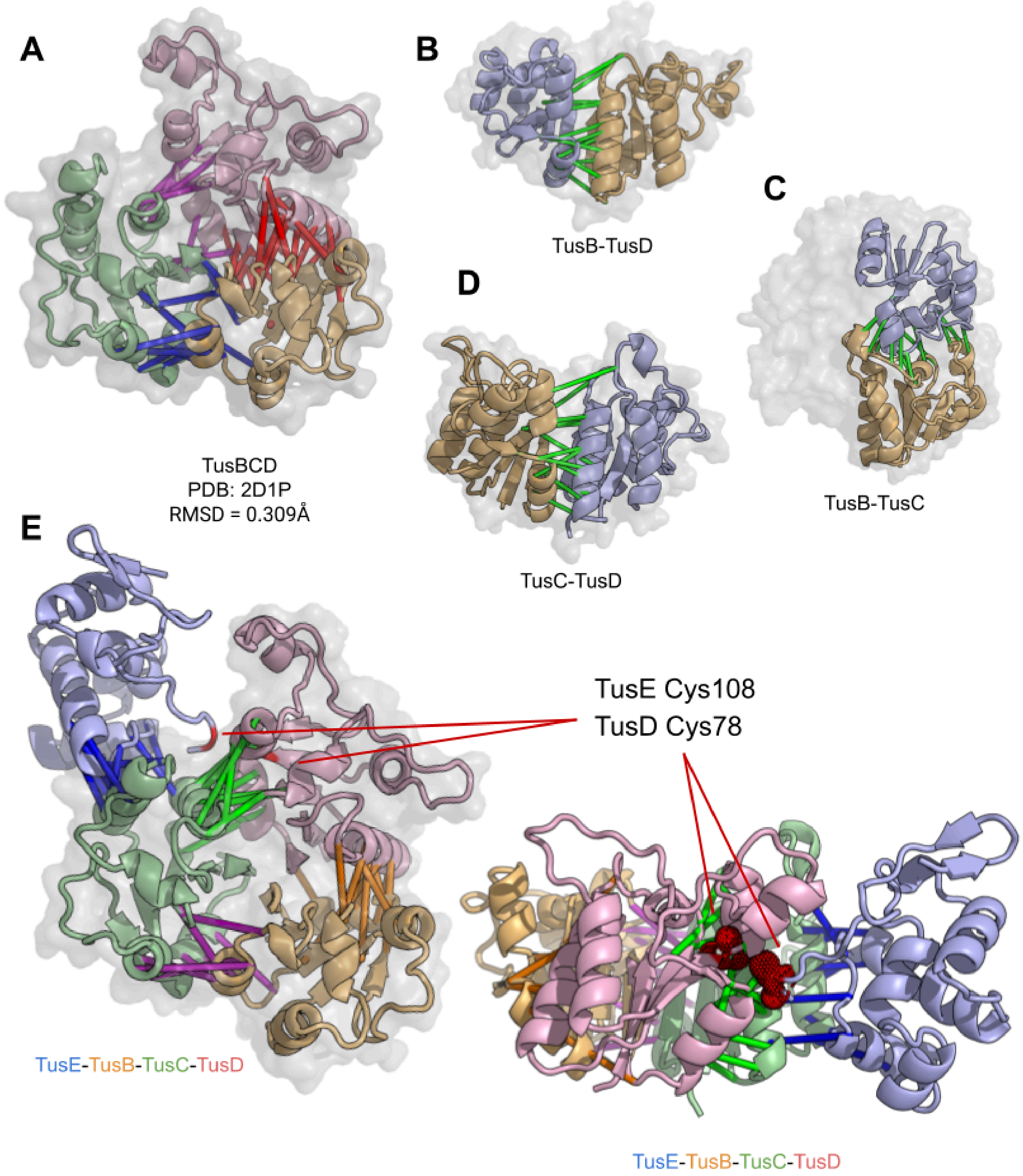


**Figure S21: tRNA 2-thiouridine synthesizing complex (Tus)**

Predictions of interactions with Tus in *E. coli*. (A-E) TusB, TusC, TusD interactions with experimentally determined TusBCD structure overlay in grey (PDB: 2D1P). (E) contains two highlighted cysteine residues which were identified as functionally relevant (*19*).

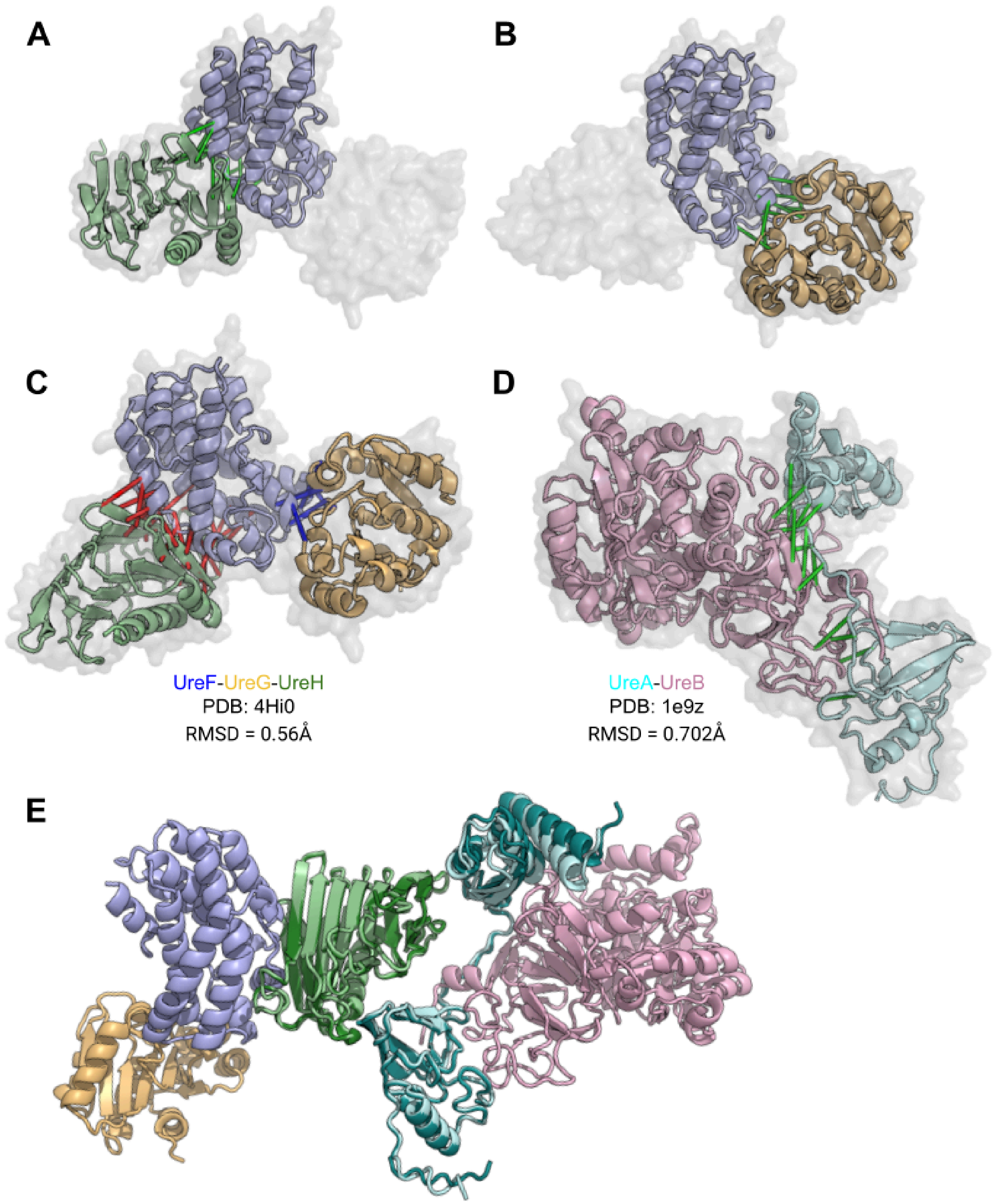


**Figure S22: Urease oligomeric assembly generation**

*H. pylori* UreAB and UreFGH complex validation through experimental structure overlays (PDB: 4Hi0, 1e9z) (*20*, *21*). (E) Pentameric UreAB-UreFGH complex assembled through multiple subcomplexes: UreFGH, UreAB, and UreAH aligned to UreAH depicted in dark teal and dark green of panel E. We note that there appear to be some steric clashes at the interface between UreA-UreH when aligned to PDB subcomplexes; however, this is not found in our predicted dimeric models. We would also like to note that we do also predict UreB-UreH, however, the interaction score from AlphaFold was 0.7, which is below our 95% precision threshold.

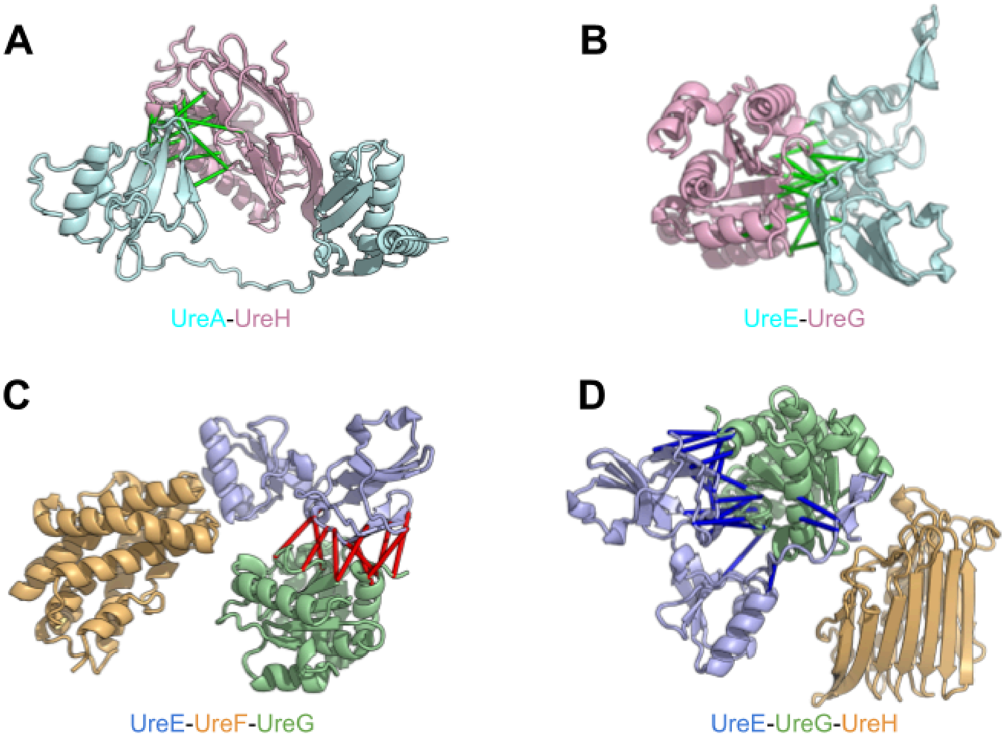


**Figure S23: Urease trimeric interactions**

Additional *H. pylori* urease predictions. (A) Dimeric interaction of UreA-UreH, which the previously described UreAB-UreFGH complex is aligned to. (B) UreE-UreG, dimeric interaction. (C,D) UreE-UreG interaction is modeled with third protein UreF or UreH, respectively, which are not predicted to interact with UreG in the presence of UreE.

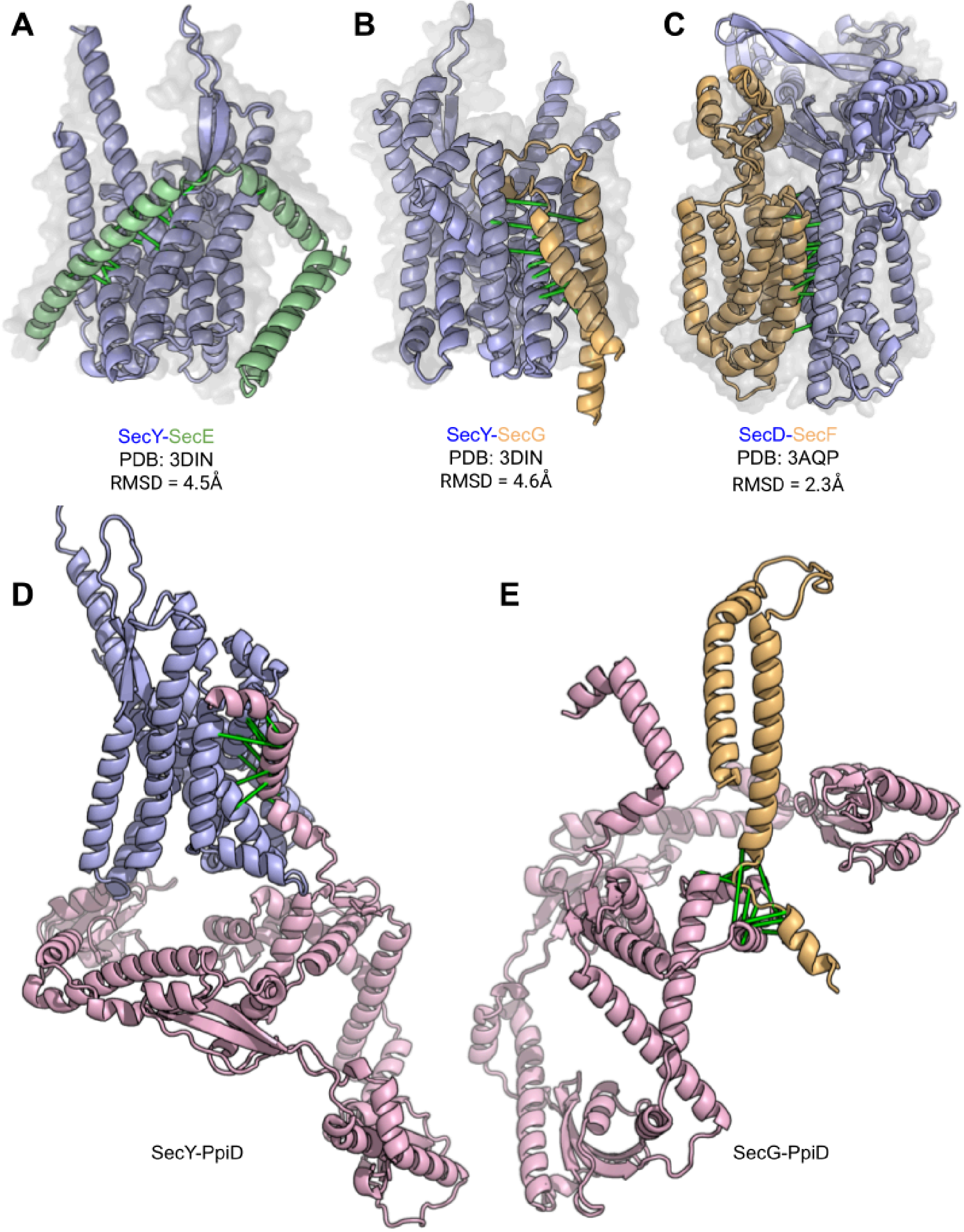


**Figure S24: Sec translocon orthologous PDB validation and PpiD**

Interactions with components of the Sec translocon in *P. aeruginosa*. (A,B) SecY-SecE and SecY-SecG dimers with overlay (PDB: 3DIN) (*22*). (C) SecD-SecF dimeric interaction with overlay (PDB: 3AQP) (*23*). (D,E) Predicted dimeric interactions with SecY/SecG and PpiD.

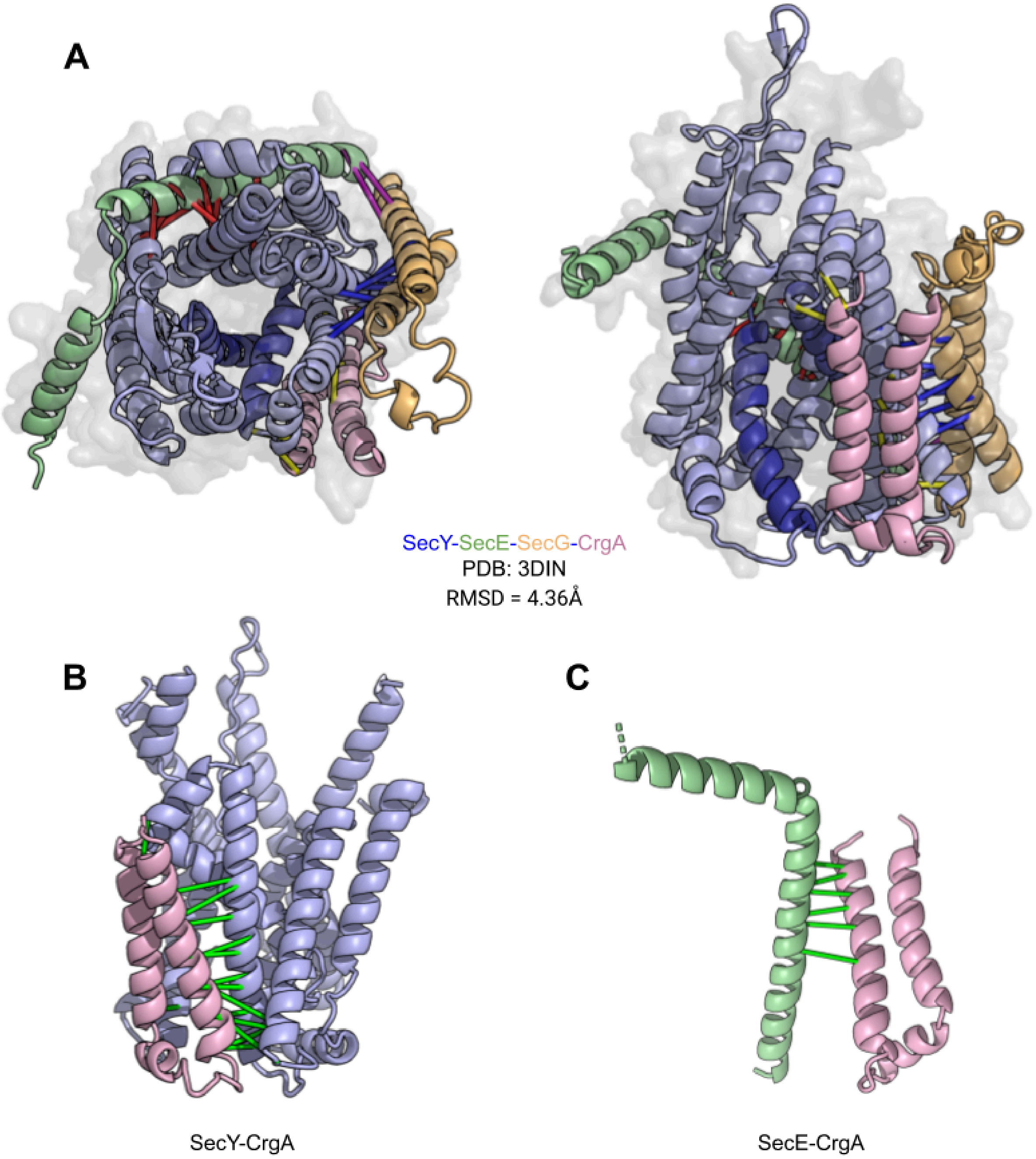


**Figure S25: Sec translocon interactions with CrgA**

Sec and CrgA interactions in *M. tuberculosis*. (A) One-shot prediction of SecYEG-CrgA with SecYEG structure overlay (PDB: 3DIN). Left top-down; right rotated 90 degrees. Transmembrane helices of two and seven colored in dark blue correspond to the lateral gate of SecY (*24*). (B,C) Dimeric interaction predictions between CrgA and Sec proteins SecY and SecE.

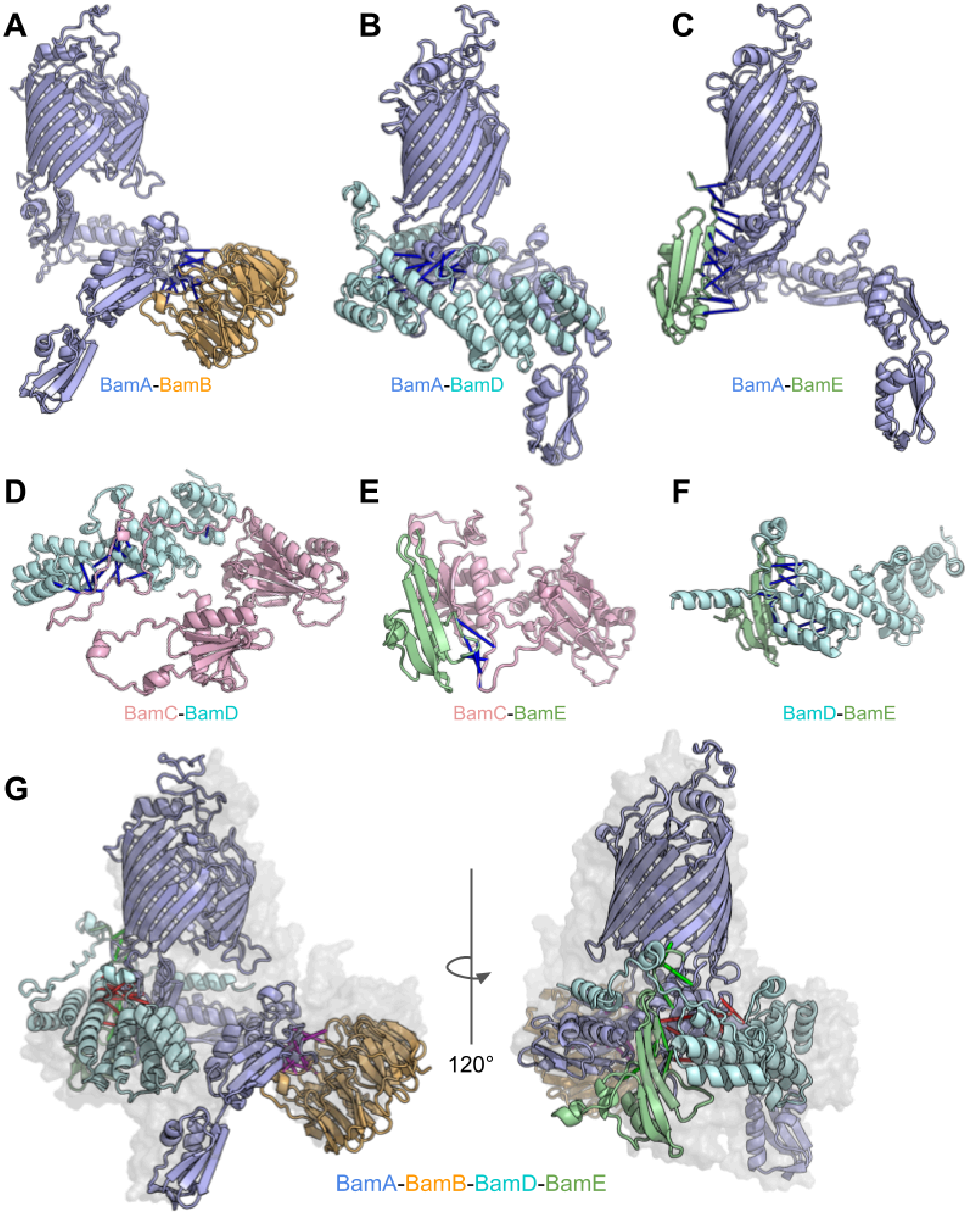


**Figure S26: BAM complex orthologous PDB validation**

Predicted interactions within the *P. aeruginosa* BamABDE complex. (A-F) predicted dimeric interactions of BamABDE. (G) Predicted one-shot tetrameric BamABDE complex with structure overlay (PDB: 5D0O) (*25*).

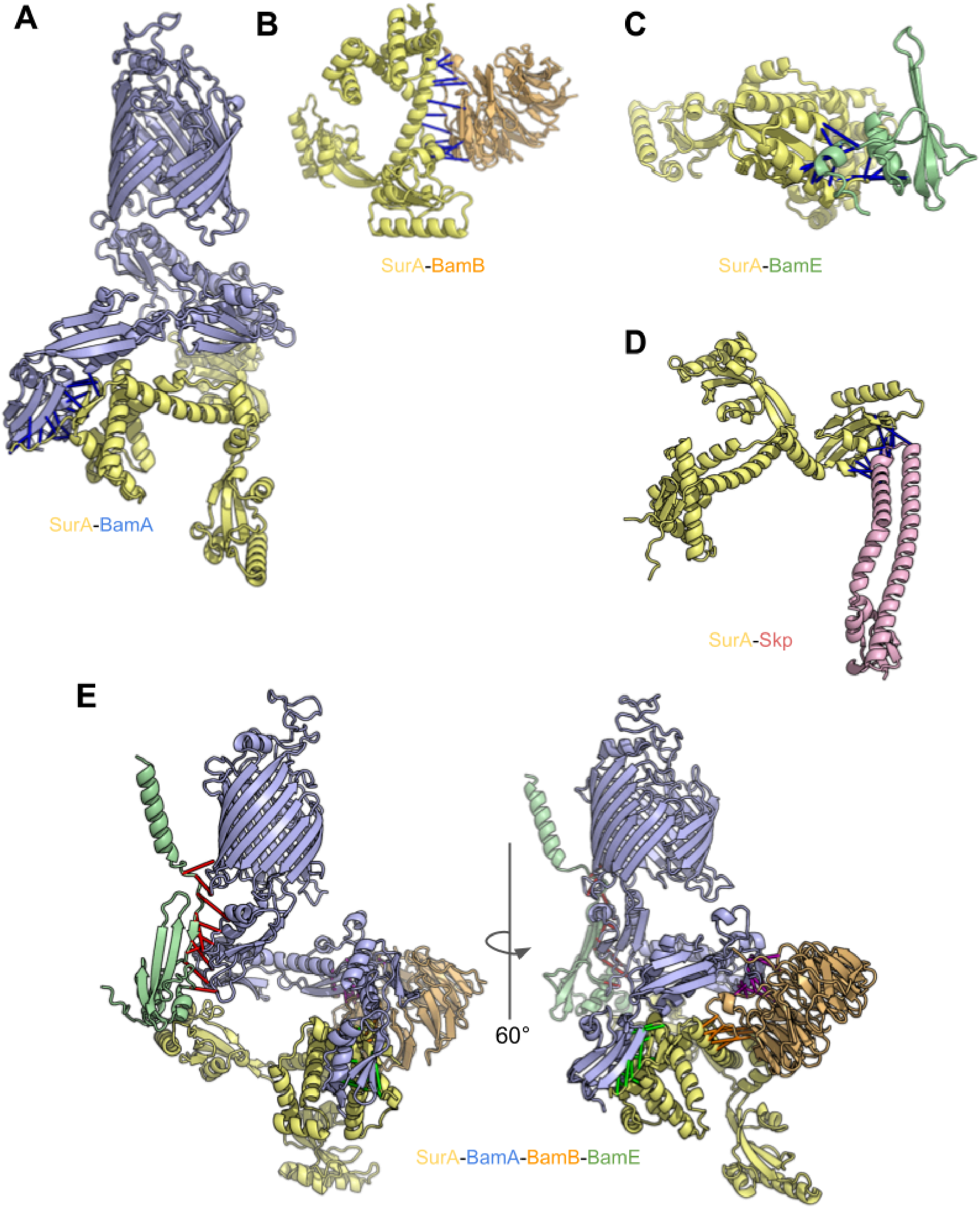


**Figure S27: BAM complex and SurA interactions**

Predicted interactions with *V. cholera* and *P. aeruginosa* Bam and SurA. (A-C) predicted dimeric interaction with SurA. (D) Predicted dimeric interaction between SurA-Skp. (E) Predicted one-shot tetrameric BamABE-SurA complex.

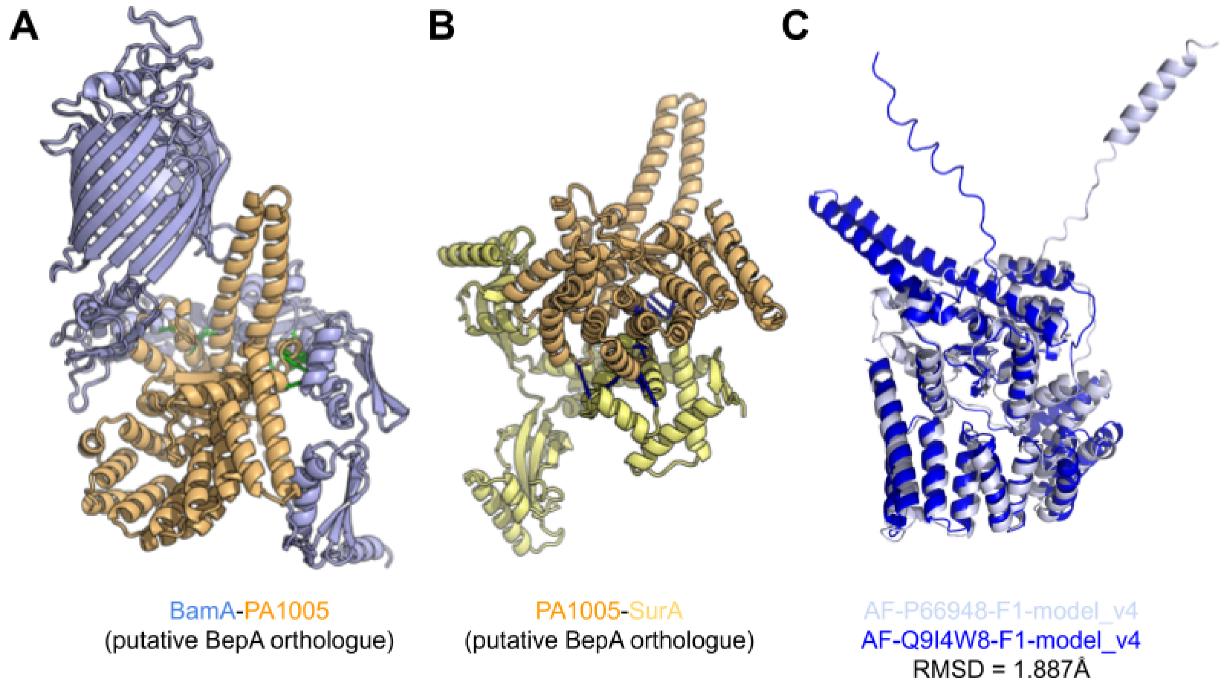


**Figure S28: BepA putative orthologue identification and Bam/SurA interaction**

Two interactions and monomeric structure of PA1005, a *P. aeruginosa* putative orthologue of BepA. (A) Predicted interaction between BamA-PA1005. (B) Predicted interaction between PA1005-SurA. (C) AlphaFold database V4 model of *P. aeruginosa* PA1005 (accession: Q9I4W8) in dark blue aligned to *E.coli* BepA (accession: AF-P66948) in light blue.

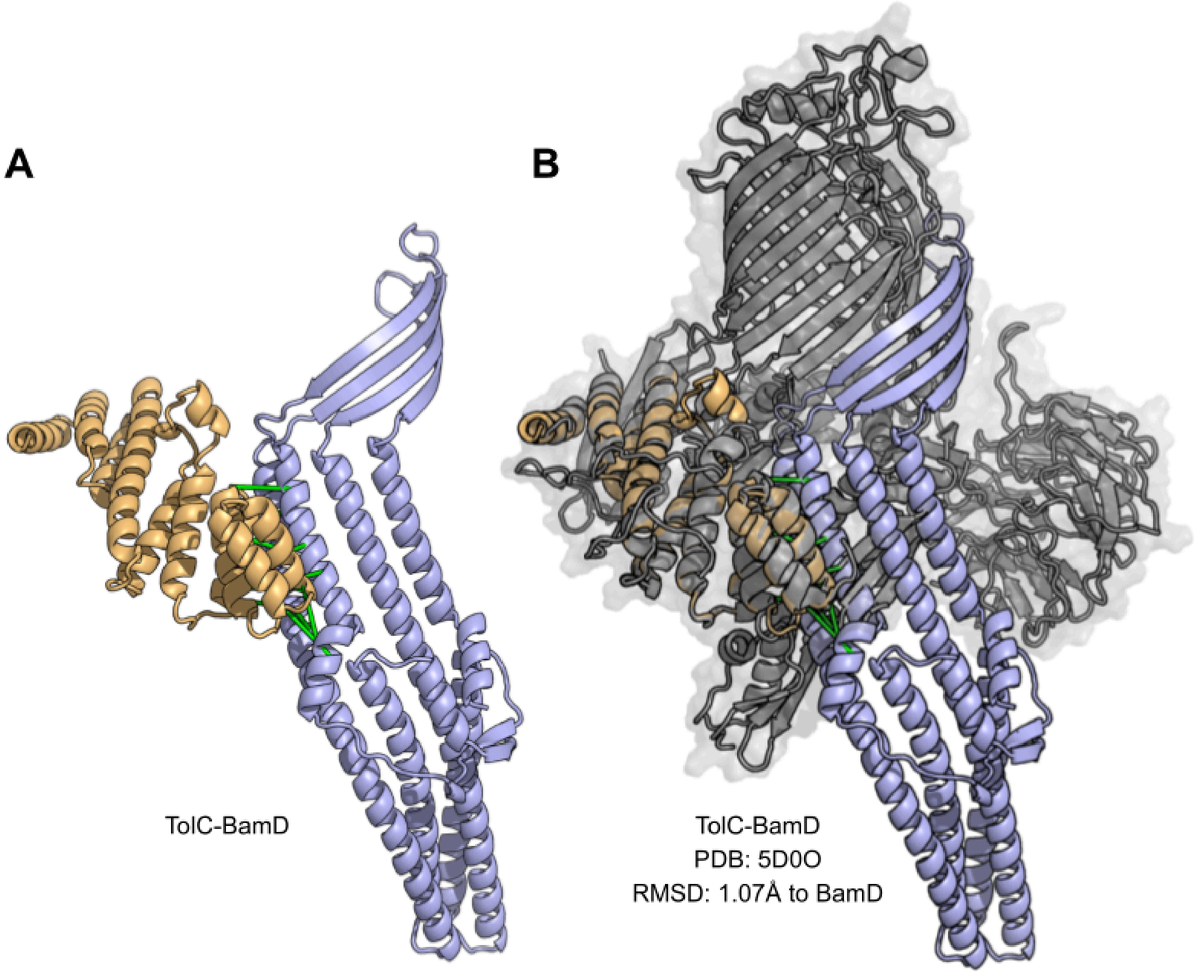


**Figure S29: Folding of TolC by BAM complex**

TolC interaction with BAM in *S. typhimurium*. (A) Predicted dimeric interaction of TolC-BamD. (B) Dimeric prediction of TolC-BamD aligned to BamD in a structure of BamABCDE complex (PDB: 5D0O); showing how the β-sheet of BamD aligns with the seam of the BamA β-barrel that folds nascent polypeptides and insertion into the outer membrane.
